# Meningeal inflammation in multiple sclerosis induces phenotypic changes in cortical microglia that differentially associate with neurodegeneration

**DOI:** 10.1101/2020.09.03.281543

**Authors:** Lynn van Olst, Carla Rodriguez-Mogeda, Carmen Picon-Munoz, Svenja Kiljan, Rachel E. James, Alwin Kamermans, Susanne M.A. van der Pol, Lydian Knoop, Evelien Drost, Marc Franssen, Geert Schenk, Jeroen J.G. Geurts, Sandra Amor, Nicholas D. Mazarakis, Jack van Horssen, Helga E. de Vries, Richard Reynolds, Maarten E. Witte

**Affiliations:** Department of Molecular Cell Biology and Immunology, Amsterdam UMC, MS Center Amsterdam, Amsterdam Neuroscience, Amsterdam, Netherlands; Division of Neuroscience, Department of Brain Sciences, Imperial College London, Hammersmith Hospital Campus, Burlington Danes Building, Du Cane Road, London W12 0NN, UK; Department of Anatomy & Neurosciences, Amsterdam UMC, MS Center Amsterdam, Amsterdam Neuroscience, Amsterdam, Netherlands; Department of Pathology, Amsterdam UMC, MS Center Amsterdam, Amsterdam Neuroscience, Amsterdam, Netherlands; Centre for Molecular Neuropathology, LKC School of Medicine, Nanyang Technological University, Singapore, Singapore; Department of Medical Biochemistry, Amsterdam UMC, Amsterdam Cardiovascular Sciences, Amsterdam, The Netherlands

## Abstract

Meningeal inflammation strongly associates with demyelination and neuronal loss in the underlying cortex of progressive MS patients, contributing to clinical disability. However, the pathological mechanisms of meningeal inflammation-induced cortical pathology are still largely elusive. Using extensive analysis of human post-mortem tissue, we identified two distinct microglial phenotypes, termed MS1 and MS2, in the cortex of progressive MS patients. These phenotypes differed in morphology and protein expression, but both associated with inflammation of the overlying meninges. We could replicate the MS-specific microglial phenotypes in a novel *in vivo* rat model for progressive MS-like meningeal inflammation, with microglia present at 1 month post-induction resembling MS1 microglia whereas those at 2 months acquired an MS2-like phenotype. Interestingly, MS1 microglia were involved in presynaptic displacement and phagocytosis and associated with a relative sparing of neurons in the MS and animal cortex. In contrast, the presence of MS2 microglia coincided with substantial neuronal loss. Taken together, we uncovered that in response to meningeal inflammation, microglia acquire two distinct phenotypes that differentially associate with neurodegeneration in the progressive MS cortex. Our data suggests that these phenotypes occur sequentially and that microglia may lose their protective properties over time, contributing to neuronal loss.

## Introduction

Multiple sclerosis (MS) is the most common chronic neurodegenerative and neuroinflammatory disease in young adults^1^. At disease onset, the majority of MS patients present with a relapsing-remitting disease course (RRMS), and despite availability of many disease-modifying therapies most RRMS patients will eventually develop secondary progressive MS (SPMS)^2–4^. The transition into SPMS is characterized by gradual worsening of neurological disability without periods of remission and currently no efficient therapeutic options exist for the majority of patients^5^. The main cause of disease progression in SPMS is neurodegeneration, which encompasses tissue damage in the white matter (WM)^6^, and accumulating pathology in the grey matter (GM) including demyelination and loss of neurons and synapses in the cerebral cortex^7–9^. Multiple studies have shown that the degree of cortical pathology provides a better correlate for progression of clinical disability than the amount of WM lesions^10–12^. The lack of therapies for SPMS can therefore be largely attributed to our incomplete understanding of the mechanisms behind cortical neurodegeneration, which, in turn, is severely hindered by absence of suitable animal models. Still, a number of studies now indicated that chronic, compartmentalized inflammation of the nearby leptomeninges is likely to drive many aspects of cortical pathology^8,13^.

Meningeal inflammation in SPMS is characterized by accumulation of immune cells like B-, T- and myeloid cells, either diffusely present or in aggregates resembling tertiary lymphoid follicles^14,15^. The degree of inflammation and the presence of follicles both associate with the severity of cortical pathology, possibly by production and subsequent diffusion of pro-inflammatory cytokines into the cortex^16,17^. Indeed, in a recently developed animal model for chronic meningeal inflammation, chronic overexpression of two of these cytokines, TNFα and IFNγ, in the leptomeninges of rats was recently shown to induce meningeal inflammation, cortical demyelination and neuronal loss, thereby closely mimicking SPMS^18^.

Microglia are the brain-resident immune cells, and as such, they continuously scan their environment for structural damage or invading pathogens using their highly motile processes^19,20^. Next to that, microglia are crucial for development and maintenance of neuronal networks by facilitating synaptic plasticity^21^ and directly interacting with neuronal cell bodies to monitor and protect neuronal function^22^. Accordingly, upon brain insults, microglia actively adapt their shape and function to restore brain homeostasis^20,23–25^. Despite this, multiple lines of evidence point towards an active contribution of microglia to neurodegeneration in many chronic neuroinflammatory and neurodegenerative diseases^26,27^. In both the previously mentioned *in vivo* model for chronic MS-like meningeal inflammation and in SPMS patients, meningeal inflammation strongly associates with activation of cortical microglia^18,28^. However, it remains to be investigated whether cortical microglia augment the pro-inflammatory signal coming from the meninges and thereby contribute to neurodegeneration in SPMS cortex.

In the SPMS cortex, activated microglia have been found in close proximity to apical dendrites, neurites and neuronal soma, but whether these microglia contribute or try to salvage ongoing neurodegeneration is currently unkown^7^. To study the link between meningeal inflammation, microglial behavior and neurodegeneration in cortical SPMS, we used post-mortem brain tissue from SPMS patients and the *in vivo* model for chronic MS-like meningeal inflammation^18^. We identified two subgroups of MS patients with distinct patterns of cortical microglial morphology, density and protein expression. Interestingly, these two subgroups of MS patients also showed separate levels of local meningeal inflammation and differed in the extent of neuronal damage. Highly similar microglial phenotypes and associations with neuronal damage were observed *in vivo*, were these phenotypes existed at different time-points after the onset of progressive MS-like meningeal inflammation. Taken together, our data suggests that in progressive MS chronic meningeal inflammation time-dependently induces phenotypic changes in cortical microglia that differentially associate with cortical neurodegeneration.

## Results

### Cortical microglia acquire a different morphological appearance in progressive MS

To assess the involvement of microglia in cortical pathology in progressive MS patients, we first quantified microglial number and morphological complexity in cortical layers 1, 3 and 5/6 using confocal microscopy on IBA1^+^ stained sections of MS patients and non-neurological controls (Fig. 1a,b). While the density of IBA1^+^ microglia in MS patients was variable and not significantly different from controls (Fig. 1c), morphological analysis of a large number of individually-traced microglia revealed more ramified microglial branching in the MS cortex, as indicated by the area under the curve (AUC) of the Sholl analysis^44^ (Fig. 1d,e). This morphological change was most evident in neuronal layer 3 (Fig. 1f) and was further corroborated on a subset of MS and control cases stained for TMEM119, a microglia-specific protein (Fig. S1b-d). Further analyzes of microglial morphology revealed an increase in the peak of the AUC which displays the maximum number of intersections in the Sholl analysis, a larger reach of the branches (wingspan), and increased number and length of branches and number of junctions in MS microglia located in neuronal layer 3 (Fig. 1f and Fig. S1e). Lastly, we observed a trend towards a larger microglial soma in all analyzed cortical layers of progressive MS patients (Fig. 1f). Co-expression of P2Y12 with virtually all non-vessel associated IBA1^+^ cells excluded the presence of IBA1^+^ infiltrating myeloid cells in our analysis (Fig. S1a).

**Fig. 1.**
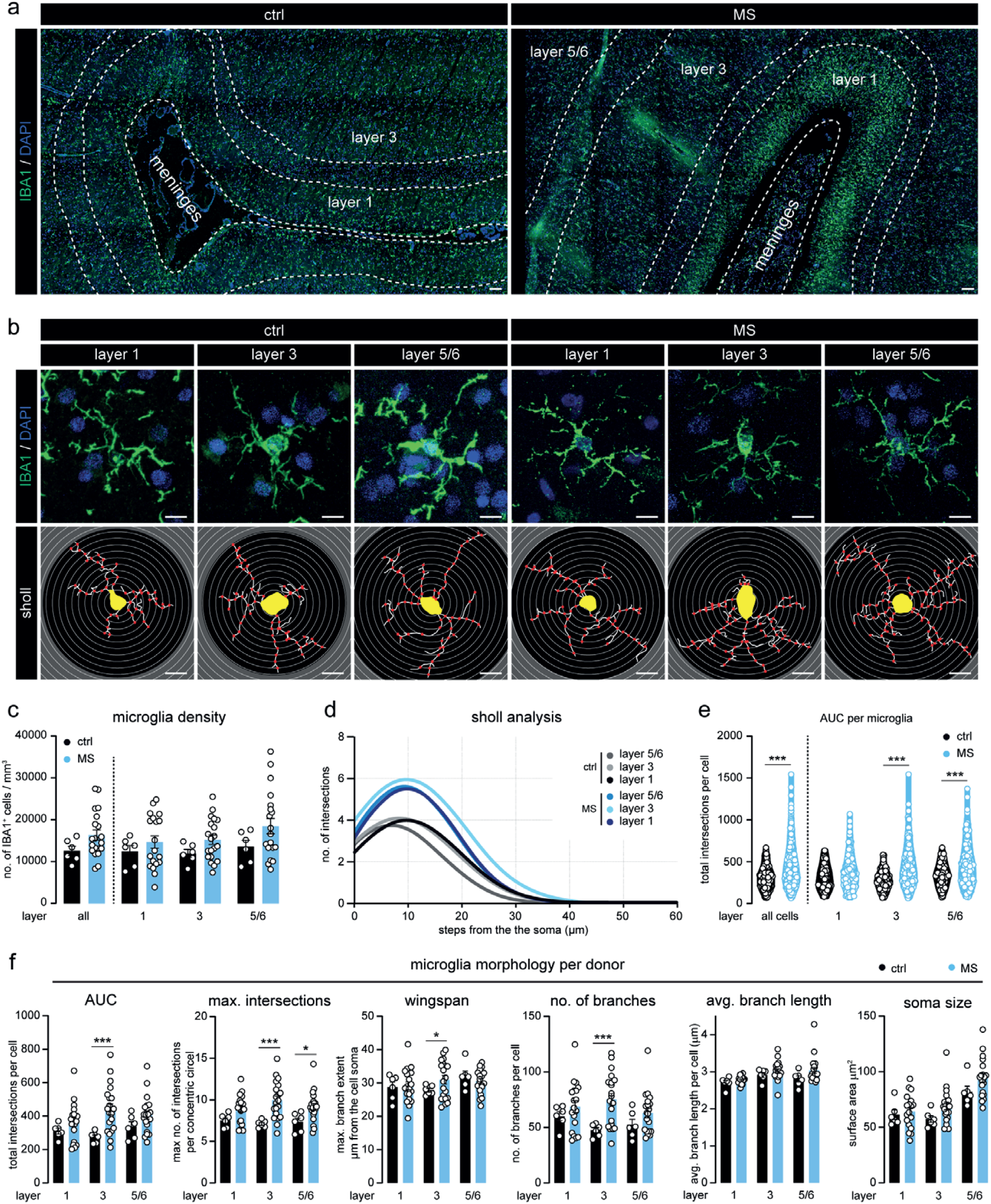
Microglia morphology is altered in MS cortex. **a**. Representative images displaying IBA1 expression in the cortex surrounding the sulcus of ctrl and MS subjects. **b**. Micrographs of individual IBA1^+^ microglia in cortical layer 1, 3 and 5/6 (top panels) and their corresponding traced outlines (bottom panels). **c**. Quantification of the microglial density per cortical layer, quantified as the number of IBA1^+^ cells per mm^3^. **d**. Non-linear curve fit of the average number of microglial branch intersections per 0.3 µm step from the cell soma per cortical layer as measured by Sholl analysis. **e**. Total Sholl-derived areaunder-the-curve (AUC) of individual microglia. **f**. Quantification of different measurements (AUC, maximal number of intersections, wingspan, number of branches, average branch length, soma size) of microglial cell morphology averaged per donor. Individual datapoints indicate averaged data from an individual donor **(c**,**f)** or individual microglia **(e)**, columns and error bars show mean ± SEM; *p < 0.05, **p < 0.01, ***p < 0.001; *n* = 6 ctrls, *n* = 20 MS subjects **(c**,**d**,**f)**, *n* = 246 microglia in ctrls, *n* = 1014 microglia in MS **(e)**; Scale bars = 10 µm **(a)**, 100 µm **(b)**.

### Heterogeneous microglial marker expression in the progressive MS cortex

To further characterize cortical microglia, we analyzed the expression of homeostatic microglia markers P2Y12 (Fig. 2a) and TMEM119 (Fig. 2b), and two commonly used activation markers, HLA class II (Fig. 2c) and CD68 (Fig. 2d), in IBA1^+^ microglia in cortical layer 3. We found that P2Y12 expression was significantly lower in MS cortical microglia compared to controls (Fig. 2b), while we observed a trend towards increased microglial HLA class II expression (Fig. 2f). TMEM119 and CD68 were not differently expressed (Fig. 2h) although expression of all 4 markers was highly variable within the MS patient group.

**Fig. 2.**
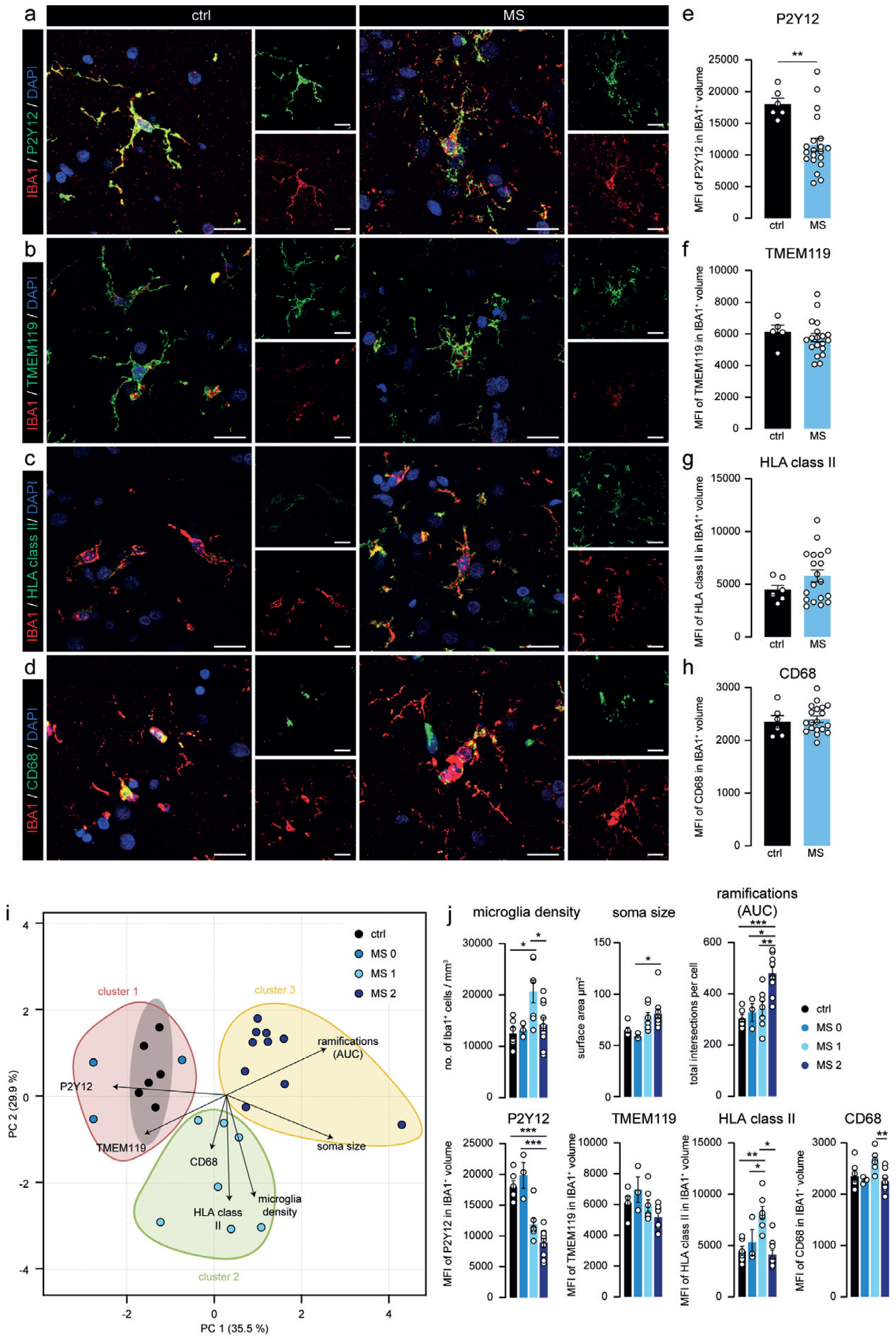
Differential expression of microglial markers in MS cortex corresponds with morphological changes. **a-d**. Representative images displaying P2Y12 **(a)**, TMEM119 **(b)**, HLA class II **(c)** and CD68 **(d)** expression in IBA1^+^ microglia in cortical layer 3 of ctrl and MS patients. **e-f**. Quantification of the mean fluorescence intensity of P2Y12 **(e)**, TMEM119 **(f)**, HLA class II **(g)** and CD68 **(h)** in microglia. **i**. Scores for the first and second principle component (PC1/PC2) of ctrl and MS patients. Subjects are coloured according to their K-means cluster assignment. **j**. Quantification of microglia density, microglia soma size, Sholl-derived area under the curve, mean fluorescence intensity of P2Y12, TMEM119, HLA class II and CD68 in IBA1^+^ volume in the different sub-groups. Individual datapoints indicate averaged data from an individual donor, columns and error bars show mean ± SEM; \**p* < 0.05, \*\**p* < 0.01, \*\*\**p* < 0.001; *n* = 6 ctrls, except in **(f)** where *n* = 5 ctrls, *n* = 20 MS subjects; Scale bars = 20 µm.

### Cortical microglia morphology, density and protein expression reveal two distinct MS subgroups

To explore if the variation in both morphology and protein expression of cortical microglia within the MS patients could be explained by the existence of subgroups of MS patients with different microglial phenotypes, we used principle component analysis (PCA) and included patient-specific parameters of microglia morphology, protein expression and cell density. PCA visualization showed the presence of three different microglial phenotypes separating MS and control cases (Fig. 2i). Then, by using K-means clustering, MS patients were assigned to three distinct subgroups which we named MS0, MS1 and MS2 (Fig. 2i). The three patients in the MS0 subgroup had a microglia phenotype that had close resemblance to those in control (Fig. 2j). MS1 was mainly defined by low P2Y12, high HLA class II, high CD68 and a markedly increased microglia density, whereas MS2 was characterized by low P2Y12 expression and increased morphological complexity as quantified by Sholl analysis (AUC), higher number of junctions and branches and increased total branch length (Fig. 2j and Fig. S2a). We did not find significant differences in age of death, disease duration, time to progressive MS and time to wheelchair between the MS subgroups (Fig. S2b). For the remainder of the study, we focused on MS-specific sub-groups MS1 and MS2, as the MS0 group was limited (n = 3) and contained microglia that were similar to those in controls.

### Increased meningeal T and B cells in progressive MS patients

Next, we explored whether meningeal inflammation is associated with the different progressive MS subgroups as several studies have already described meningeal inflammation as an important driver of cortical pathology in progressive MS^8,13,19^. We found that CD19^+^ B cells and CD3^+^ T cells were increased in the meninges of progressive MS patients, while the levels of meningeal IBA1^+^ myeloid cells remained unchanged (Fig. 3a,b and Fig. S3a). Interestingly, the increase in meningeal CD19^+^ B cells was most notable in the MS2 group (Fig. 3c), and correlated with increased morphological complexity of cortical microglia. To further dissect the involvement of meningeal T cells, we analyzed the presence of CD4^+^ and CD8^+^ T cells (Fig. 3d). As expected, both CD4^+^ and CD8^+^ T cells numbers were increased in a large subset of MS patients (Fig. 3e, and Fig. S3b), and similar CD4/CD8 T cell ratios shows that they were equally induced in meninges from patients with MS1 and MS2 microglial phenotypes. (Fig. 3f).

**Fig. 3.**
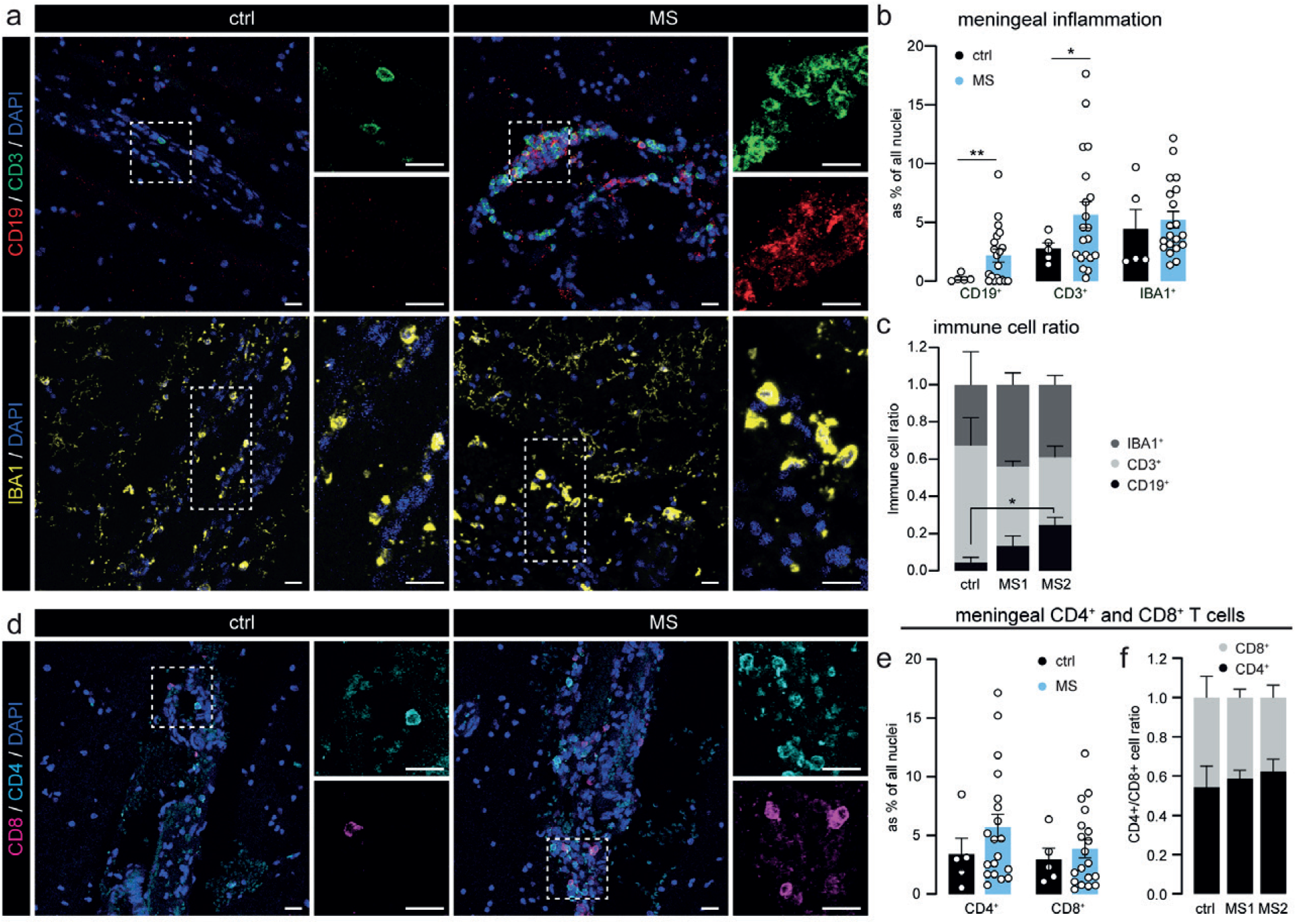
MS2 microglia associate with meningeal B cells in progressive MS. **a**. Representative images of the leptomeninges in ctrl and MS patients immunostained for CD3 (T cells), CD19 (B cells) and IBA1 (myeloid cells). **b**. Quantification of CD19^+^, CD3^+^ and IBA1^+^ cells as percentage of all nuclei in the meninges. **c**. Immune cell ratio of CD19^+^, CD3^+^ and IBA1^+^ cells in the meninges of ctrls, MS1 and MS2 clusters. **d**. Representative images of CD8^+^ (pink) and CD4^+^ (cyan) T cells in meninges of ctrl and MS patients. **e**. Quantification of CD4^+^ and CD8^+^ cells as percentage of all nuclei in the meninges. **f**. CD4^+^/CD8^+^ immune cell ratio in the meninges of ctrls, MS1 and MS2 clusters. Individual datapoints indicate averaged data from an individual donor, columns and error bars show mean ± SEM; *p < 0.05, **p < 0.01; *n* = 5 ctrls and *n* = 20 MS subjects **(b)**, *n* = 3 ctrls, *n* = 5 MS1 and *n* = 6 MS2 **(c)**, *n* = 5 ctrls and *n* = 19 MS subjects **(e)**, *n* = 4 ctrls, *n* = 7 MS1 and *n* = 9 MS2 **(f)**; Scale bars = 20 µm.

### Experimental chronic meningeal inflammation time-dependently induces MS1- and MS2-like microglia

To investigate whether meningeal inflammation could drive the microglial changes we observed in the progressive MS cortex, we made use of a novel animal model of chronic experimental meningeal inflammation (CMI; Fig. S4a) which has recently been shown to replicate important features of cortical pathology in progressive MS patients^18^. In this model, a sub-clinical autoimmune response to myelin was induced and lentiviral vectors containing TNFα and IFNγ were injected in the sagittal sulcus below the meningeal dura mater layer. As expected, we observed an increase in meningeal CD3^+^ T cells, IBA1^+^ myeloid cells and CD79a^+^ B cells in the sagittal sulcus of CMI rats 1 and 2 months after lentiviral injection. Especially CD79a^+^ B cells were most strongly increased in number after 2 months (Fig. 4a-c and Fig. S4b). Similar to the previous study, we did not observe significant differences in meningeal immune cell infiltration between CMI rats, which underwent subclinical MOG immunization prior to lentiviral injection, and IFA control rats, which were only injected with IFA prior to lentiviral injections (Fig. S4c).

**Fig. 4.**
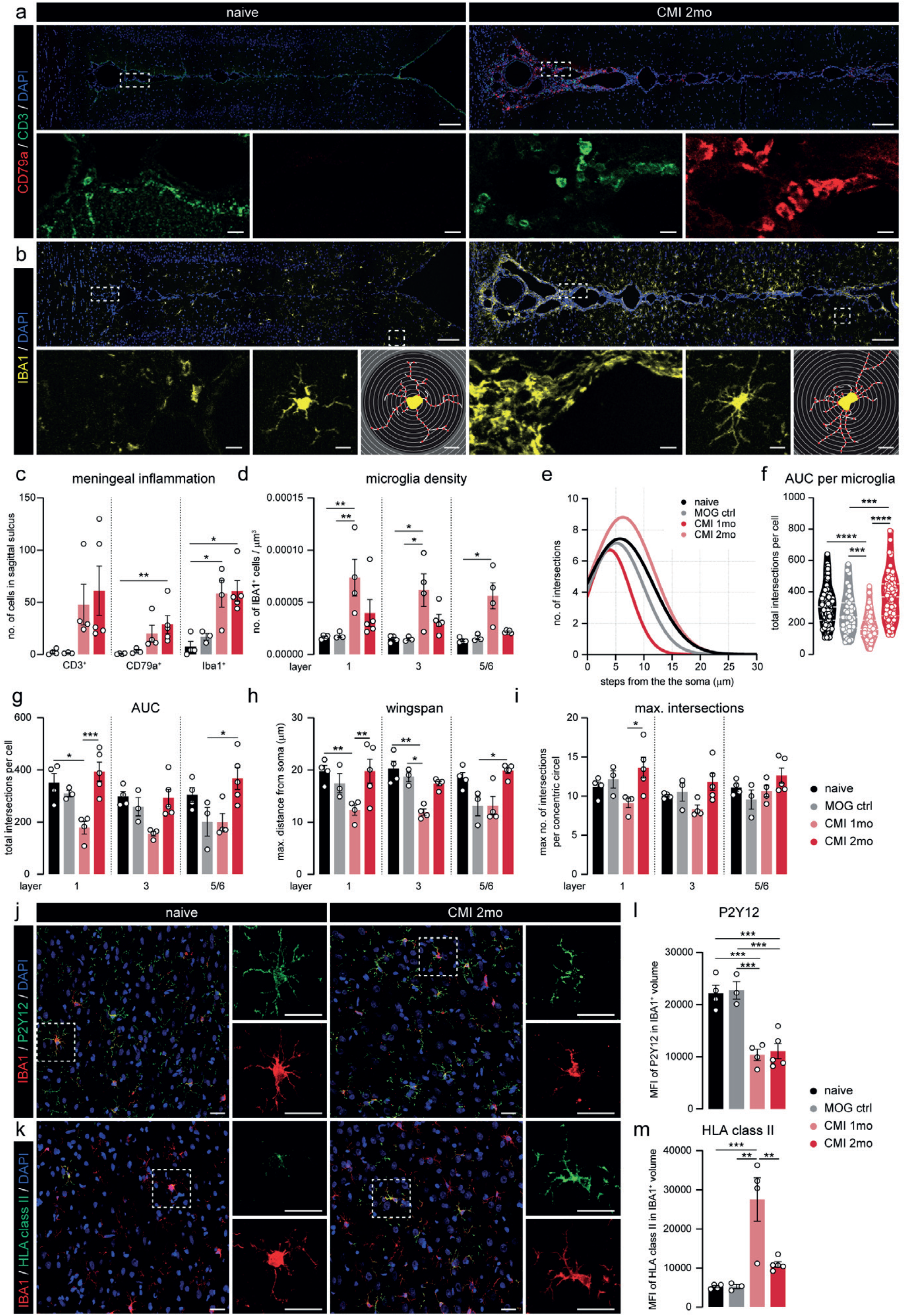
Experimental chronic meningeal inflammation induces similar microglial phenotypes as in MS cortex at different time points. **a**. Representative image of CD3 (T cells) and CD79a (B cells) expression of the sagittal sulcus and surrounding cortex of naive and CMI 2 months animals. **b**. Representative images of IBA1 expression in and around the sagittal sulcus of naïve and CMI 2 month rats (top panels). Higher magnification images of IBA1 expression inside the meninges (lower panel - left). Close-up of a single IBA1^+^ cell (lower panel - middle) and corresponding traced outline (lower panel - right). **c**. Absolute number of CD3^+^ T cells, CD79a^+^ B cells and IBA1^+^ cells in the sagittal sulcus. **d**. Microglial density per cortical layer, quantified as the number of IBA1^+^ cells per µm^3^ in the different animal groups **e**. Non-linear curve fit of the average number of microglial branch intersections per 0.3 µm step from the cell soma per cortical layer as measured by the Sholl analysis. **f**. Total Sholl-derived area-under-the-curve (AUC) of individual microglia. **g-i**. Different measurements (AUC, wingspan, maximal number of intersections) of microglial cell morphology averaged per animal. **j, k**. Representative confocal images of cortical layer 3 from naïve and CMI 2 months displaying P2Y12 **(j)** or HLA class II **(k)** and IBA1 expression. **i, m**. Mean fluorescence intensity of P2Y12 signal **(i)** and HLA class II **(m)** in IBA1+ volume. Individual datapoints indicate averaged data from an individual donor **(c-e**,**g-i**,**l**,**m)** or individual microglia **(e)**, columns and error bars show mean ± SEM; *p < 0.05, **p < 0.01, ***p < 0.001; *n* = 4 naïve, *n* = 3 MOG ctrl, *n* = 4 CMI 1mo, *n* = 5 CMI 2mo **(c-e**,**g-i**,**l**,**m)**, *n* = 112 naïve, *n* = 43 MOG ctrl, *n* = 173 CMI 1mo, *n* = 143 CMI 2mo **(f)**; Scale bars = 100 µm (top panels of **a**,**b**), 10 µm (lower panels of **a**,**b)**, 25 µm **(j**,**k)**.

Next, we compared microglial density, morphology and protein expression in the cortex surrounding the sagittal sulcus in CMI rats with appropriate controls. Here, we found that after 1 month, CMI animals displayed an increased microglia density (Fig. 4d) and a less ramified morphology (Fig. 4e-i). In contrast, after 2 months, microglial density was almost back to control levels (Fig. 4b-d) whereas morphological complexity of microglia was similar to or greater than in controls (Fig. 4e-f). We observed little to no changes in microglial soma size in both 1 and 2 month CMI animals (Fig S4d). Importantly, virtually all non-vessel associated IBA1^+^ cells in the cortex of CMI rats were positive for P2Y12, indicating very little or absent infiltration of peripheral myeloid cells (Fig. S4e). Quantification of P2Y12 and HLA class II expression in cortical layer 3 microglia of CMI rats (Fig. 4j-m) revealed a clear reduction in P2Y12 expression after both 1 and 2 months (Fig. 4j-i), whereas expression of HLA class II was highly upregulated in CMI 1 month microglia but almost back to levels seen in control animals after 2 months (Fig. 4m). Again, we found similar changes in cortical microglial phenotype in IFA control animals after 1 and 2 months, albeit that most changes were less pronounced than in the MOG-immunized CMI animals (Fig. S4f).

Taken together, microglia in CMI rats differ substantially between 1 month from those 2 months after lentiviral injection. Interestingly, those seen after 1 month carry many features of MS patients previously localized to the MS1 subgroup, including a high microglial density, a less ramified morphology, low P2Y12 and high HLA class II expression. In addition, characteristics of CMI rats 2 months after injection, i.e. higher number of meningeal B cells, lower microglial density and HLA class II expression, but an increased morphological complexity, closely resembles the MS2 subgroup.

### MS1 and MS2 subgroups differentially associate with cortical neurodegeneration

We next questioned whether the two MS clusters differentially associate with local tissue damage. For this purpose, we first quantified demyelination in MOG-stained sections and, as expected, we observed that most progressive MS patients had one or more cortical lesions, mostly of the subpial type, which were completely absent from controls (Fig. 5a). However, we did not detect any differences between MS1 and MS2 patients in either lesion load (as % of total cortex; Fig. 5b) or MOG^+^ area in the different cortical layers (Fig. 5c). We subsequently compared neuronal density per cortical layer using HuC/D immunolabeling (Fig. 5d). As it was recently reported that neurons in layer 2 and 3 of the cortex are especially prone to degenerate in progressive MS cortex^29^, we also decided to include cortical layers 2 and 4 in this analysis. In line with a previous study^8^, we observe a lower neuronal density in the upper cortical layers of most progressive MS patients, but this only reached statistical significance in cortical layers 2 and 3 of MS2 patients (Fig. 5e).

**Fig. 5.**
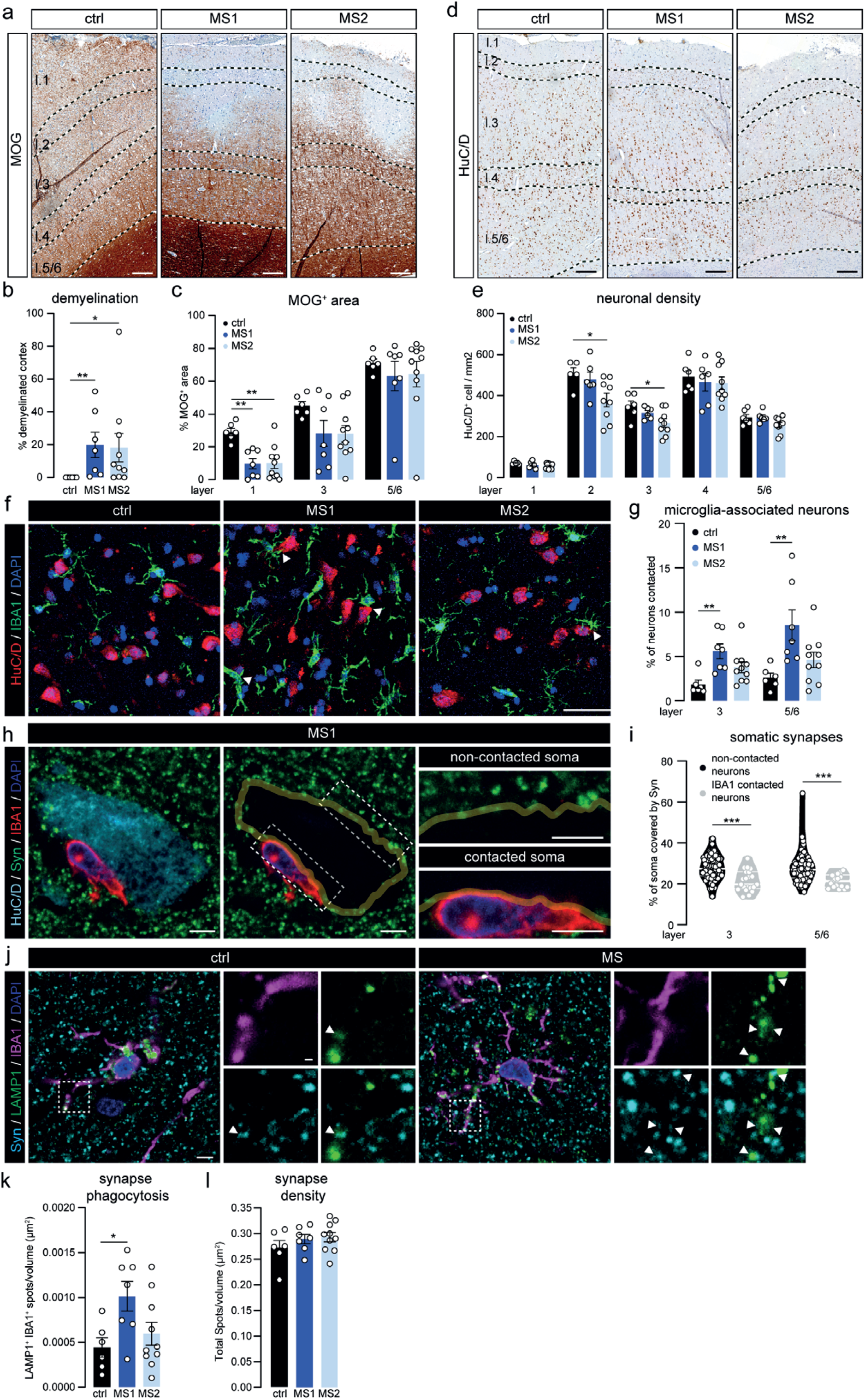
MS subgroups based on microglia phenotype reveal different levels of neuronal loss and synaptic removal. **a**. Representative images of MOG-stained cortex of ctrl, MS1 and MS2 subjects. **b-c**. Percentage of total cortical demyelination **(b)** and MOG^+^ pixels **(c)** in cortical layers 1, 3 and 5/6 d. Representative images showing HuC/D-labeled neurons in the cortex of ctrl, MS1 and MS2 subjects. **e**. Quantification of neuronal density in the different cortical layers. **f**. Representative images of cortical layers 5/6 from a ctrl, MS1 and MS2 case double-labeled with IBA1 (microglia) and HuC/D (neurons). Arrow heads depict soma-soma contact between microglia and neurons. **g**. Percentage of neuronal somata directly contacted by microglia in cortical layers 3 and 5/6. **h**. Representative single z-plane of a cortical layer 3 neuron from an MS1 subject immunolabeled for HuC/D (neuronal soma), IBA1 (microglia) and Synaptophysin (pre-synapse) (left panel). Yellow line depicts the outline of the neuronal cell body (middle panel), highlighting the synaptic input on the soma. Right panels show close-ups of the areas outlined in the middle panel. **i**. Quantification of the percentage of neuronal soma covered by Synaptophysin^+^ structures in microglia-contacted neurons and neurons not associated with microglia in the MS cortex (pooled data from 3 MS1, and 3 MS2 cases). **j**. Representative images of cortical layer 3 from a ctrl and MS1 subject stained for IBA1 (microglia), LAMP1 (lysosomes) and Synaptophysin (pre-synapses). **k**. Number of Synaptophysin^+^ spots in microglial lysosomes per volume of cortical layer 3. **l**. Total Synaptophysin^+^ spot density in cortical layer 3. Individual datapoints indicate averaged data from an individual donor **(b**,**c**,**e**,**g**,**k**,**l)**, or individual neurons **(i)**, columns and error bars show mean ± SEM; *p < 0.05, **p < 0.01, *** p < 0.001; *n* = 6 ctrls **(b**,**c**,**e**,**g**,**k**,**l)** except in layer 1 **(c**,**e)** and layer 2 **(e)** where *n* = 5 ctrls, *n* = 7 MS1 and *n* = 10 MS2 **(b**,**c**,**g**,**k**,**l)**, *n* = 6 MS1 and *n* = 9 MS2 **(e)**, *n* = 110 layer 3 and *n* = 102 layer 5/6 neurons **(i)** ; Scale bars = 250 µm **(a**,**d)**, 50 µm **(f)**, 10 µm **(h**,**j)**.

Several studies have described close contact between microglia and neurons in neuro-inflammatory conditions^24,30^, leading to removal of pre-synaptic input from the neuronal soma. Here, using microglial IBA1 and neuronal HuC/D co-labeling, we show that in the cortex of MS1 patients a higher percentage of neuronal somata is directly associated with one or more microglial cell bodies (Fig. 5f-g). Similarly, we also observe a higher percentage of microglia that are contacting neurons in both MS1 and MS2, indicating that the increased neuron-microglia association is not just a bystander effect of increased microglial density in progressive MS (Fig. S5a). Next, we found an almost complete loss of Synaptophysin^+^ (Syn) pre-synaptic input on that part of the neuronal soma which was occupied by a microglial cell body (Fig. 5h). In addition, we show an overall reduction of pre-synaptic input on neuronal soma in direct contact with one or more microglia in MS cortex (Fig. 5i).

Next, we assessed whether cortical microglia in MS1 and MS2 patients were engaged in phagocytosis of pre-synapses by quantifying the presence of Syn^+^ structures in LAMP1^+^ lysosomes in microglia (Fig. 5j). Similar to what happens in the thalamus of MS patients^31^, we found a significant increase in synaptic phagocytosis specifically in MS1 cortical microglia (Fig. 5j-k), which is in line with an overall increased phagocytic capacity of MS1 microglia as indicated by increased CD68 expression (Fig. 2) and corroborated here by increased LAMP1 expression (Fig. S5b). Remarkably, we did not observe a decrease in the total pre-synaptic density in layer 3 of either MS1 or MS2 patients (Fig 5l), which might be explained by the fact that the percentage of all synapses that were located in microglial lysosomes was only around 0.3% (Fig. S5c). A small fraction of synapses where located in lysosomes of non-microglial cells, but there was no difference between control, MS1 and MS2 donors (Fig. S5d).

### CMI induces time-dependent cortical neurodegeneration

To assess whether we find similar associations with cortical tissue damage in CMI animals as we did in progressive MS patients, we first measured demyelination in the cortical layers extending from the sagittal sulcus in MOG-stained sections (Fig. 6a). As shown previously, CMI induces cortical demyelination most strongly in CMI 2 months and in the layers closest to the sulcus^18^. Due to the large variability of MOG^+^ area in all groups, we did not detect any significant difference in layers 1 or 2 (Fig. 6b). Moreover, neuronal density in layer 1 was decreased at both time points, whereas we only found neuronal loss in layer 2 after 2 months (Fig. 6c-d). Remarkably, we also observed a small but significant loss of NeuN^+^ neurons in layers 4-6 after 1 month, but not after 2 months (Fig. 6d).

**Fig. 6.**
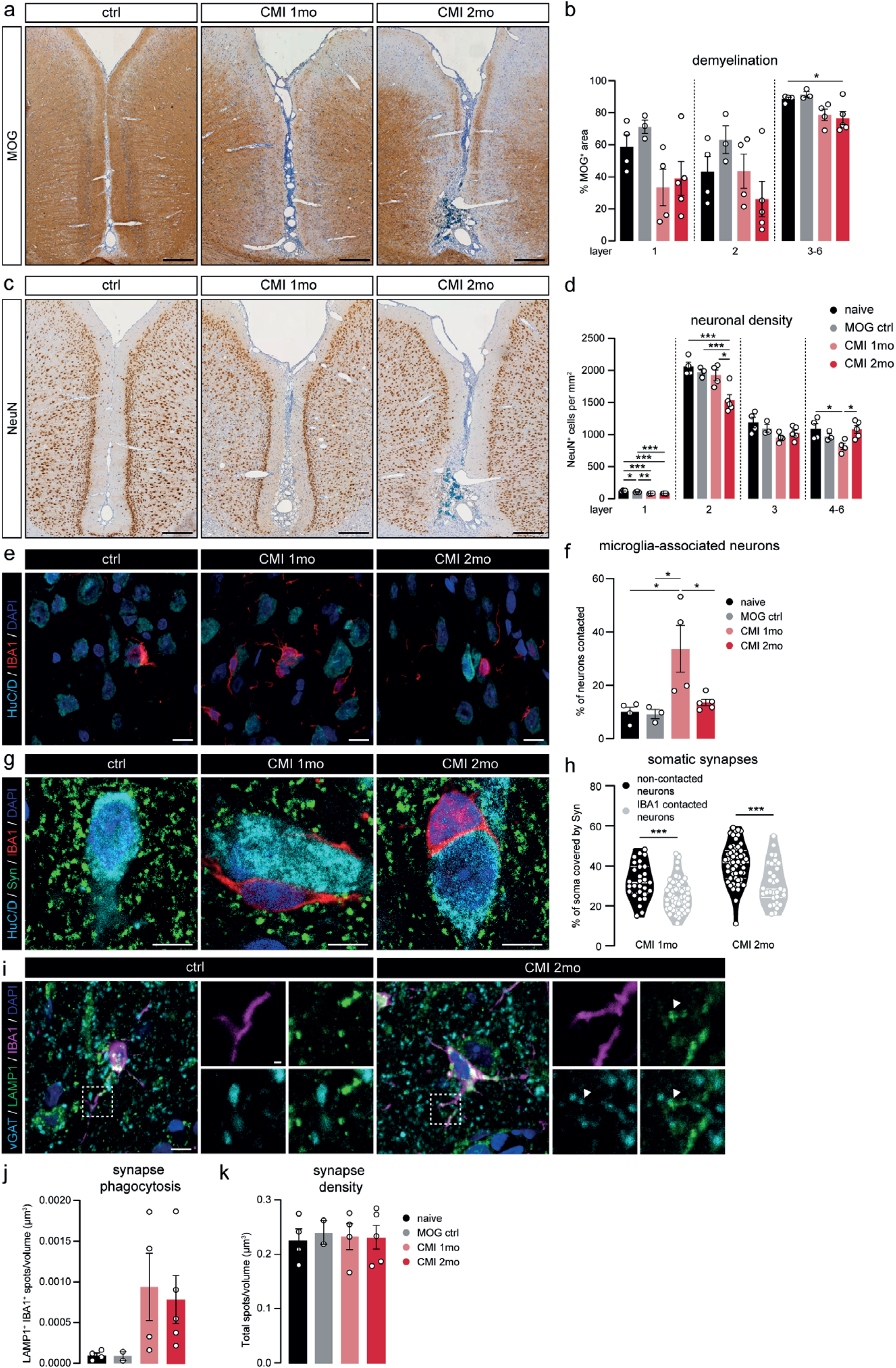
Experimentally induced chronic meningeal inflammation recapitulates cortical pathology in the MS cortex. **a**. Representative images of MOG-stained cortex around the sagittal sulcus of naive, CMI 1 month and CMI 2 months animals. **b**. Quantification of the percentage of MOG^+^ pixels in cortical layers 1, 2 and 3-6 surrounding the sagittal sulcus in the four groups of rats. **c**. Representative images showing NeuN^+^ neurons in the cortex of naive, CMI 1 month and CMI 2 months animals. **d**. Quantification of neuronal density in the different cortical layers. **e**. Representative maximum projection confocal images of cortical layer 3 from naive, CMI 1 month and CMI 2 months animals double-labeled with IBA1 (microglia) and HuC/D (neurons). **f**. Quantification of the percentage of neuronal somata directly in contact with microglia soma in cortical layer 3. **g**. Representative single z-plane confocal images of cortical layer 3 of naive, CMI 1 month and CMI 2 months animals immunolabeled for HuC/D (neuronal somata), IBA1 (microglia) and Synaptophysin (pre-synapses). **h**. Quantification of the percentage of neuronal soma covered by Synaptophysin-labeled structures in microglia-contacted neurons and neurons not associated with microglia in CMI animals (pooled data from 3 animals in each group). **i**. Representative images of IBA1 (microglia), LAMP1 (lysosomes) and vGAT (pre-synapses) expression in cortical layer 3 of a naive and CMI 2 months animal. **j**. Quantification of the density of vGAT^+^ spots in microglial lysosomes of cortical layer 3. **k**. Total density of vGAT spots in layer 3 of the cortex. Individual datapoints indicate averaged data from an individual animal **(b**,**d**,**f**,**h**,**j**,**k)**, or individual neurons **(h)**, columns and error bars show mean ± SEM; *p < 0.05, **p < 0.01, ***p < 0.001; *n* = 4 naïve, *n* = 3 MOG ctrl, *n* = 4 CMI 1mo, *n* = 5 CMI 2mo **(b**,**d**,**f**,**h**,**j**,**k)** with the exception of *n* = 2 MOG ctrl in **(j**,**k)**, *n* = 80 CMI 1mo and *n* = 88 CMI 2mo neurons **(h)**,. Scale bars = 250 µm **(a**,**c)**, 10 µm **(e**,**g)**.

In line with what we observe in the cortex of MS1 cases, the number of neuronal somata directly associated with microglia cell bodies was significantly increased in layer 3 of CMI 1 month rats, whereas after 2 months this was almost back to control levels (Fig. 6e-f). At both time points, these contacts resulted in removal of pre-synaptic input from the neuronal soma (Fig. 6g-h). Phagocytosis of vGAT^+^ pre-synapses by microglia was increased after 1 and 2 months, albeit with a large variation between animals (Fig. 6i-j). Again, we did not observe a total loss of vGAT^+^ synapses in cortical layer 3 of CMI rats (Fig. 6k), possibly due to the small percentage of pre-synapses found in microglial lysosomes (Fig. S6a). Furthermore, synapse phagocytosis by non-microglia cells was not significantly different between groups (Fig. S6b).

## Discussion

Currently, therapeutic options for progressive MS patients are limited^5^. This is partly attributed to our lack of understanding of the pathological mechanisms driving the disease, which in turn is a consequence of a dearth of suitable animal models for progressive MS^32^. In this study, using both post-mortem tissue from progressive MS patients and a recently developed animal model for progressive MS-related cortical pathology^18^, we uncover mechanisms that drive neurodegeneration in progressive MS cortex and further establish the use of this new animal model to study underlying pathological processes.

By using extensive morphological analysis and quantification of several well-known markers for microglial activation, we could separate our cohort of progressive MS patient tissue into three distinct clusters. Two of these clusters (termed MS1 & MS2) contained a cortical microglia population that was significantly different from controls. MS1 microglia were characterized by a high cellular density and elevated expression of both HLA class II and CD68; whereas MS2 microglia were defined by a hyper-ramified morphology and low P2Y12 expression. Both MS-specific microglia phenotypes were linked to increased meningeal inflammation, but only MS2 associated with the presence of meningeal B cells. Remarkably, we could detect microglia that were very similar to both MS1 and MS2 in an animal model for chronic MS-like meningeal inflammation (CMI). Cortical microglia found 1 month after the induction of meningeal inflammation resembled MS1 microglia, while microglia at 2 months after induction shared many features with MS2 microglia. We further show that MS1 microglia spatially associated with neuronal cell bodies, whereas significant neuronal loss was restricted to patients assigned to the MS2 cluster. The spatial association between microglia and neurons was related to the removal of pre-synapses from the neuronal soma, and accompanied by increased pre-synapse phagocytosis by MS1 microglia. Again, neuronal changes in CMI animals reflected what we observed in progressive MS tissue, with cortical microglia 1 month after induction of meningeal inflammation closely apposing neuronal somata, and neuronal loss in the upper cortical layers being most prominent after 2 months. Furthermore, we observed a removal of pre-synapses from the soma of neurons that were contacted by microglia in CMI animals and found evidence for synapse phagocytosis by microglia at both time-points.

Given the striking similarities between the two MS-specific clusters and the cortex of CMI animals after 1 and 2 months, it is tempting to speculate that the changes observed in MS1 cortex are caused by early stage meningeal inflammation and those observed in MS2 cortex by more chronic meningeal inflammation. As B cells are most prominent in the meninges of both MS2 patients and 2 month CMI animals, this would suggest that at first T cells and myeloid cells populate the MS meninges, with the amount of meningeal B cells rising over time. This also fits with the finding that the amount of B cells in meninges and B-cell related cytokines in the CSF associate with disease progression^17,28,33^. Furthermore, the extent of meningeal inflammation in MS patients, and more specifically the amount of meningeal B cells, has been strongly linked to the presence of tertiary lymphoid-like follicles^15,28^. Unfortunately, we could identify too few of these lymphocyte clusters in the meninges of our tissue cohort to test whether they indeed are found predominantly in the MS2 cluster, as one would expect. Likewise, it would be interesting to explore whether these follicles also occur in CMI animals, which could open up studies examining their development and, eventually, used to find therapeutics that could block their formation.

Microglia have been extensively studied in the context of age-related neurodegenerative diseases and white matter lesions in MS (for review see^26,27^), however their role in MS-related cortical pathology remains relatively obscure. Studies in amyotrophic lateral sclerosis, Alzheimer’s disease and white matter MS lesions and their respective animal models have identified what appears to be a shared microglial response to (neuronal) damage^34^, which involves retraction of their processes and significant changes in protein and/or mRNA expression. This phenotype has been termed either MGnD (microglia neurodegenerative phenotype) or DAM (disease-associated microglia)^35^, but whether these microglia are beneficial or detrimental remains a matter of debate and could very well be time and disease-dependent. Here, we show that cortical microglia in MS respond rather differently and acquire a more ramified morphology throughout the cortex, especially in the MS2 cluster. Similarly, we found that markers known to be upregulated under neuroinflammatory and neurodegenerative conditions, HLA and CD68^36,37^, are not altered in MS2 microglia. Lastly, the expression of the homeostatic marker TMEM119 did not significantly differ from controls in all clusters unlike in MGnD/DAM microglia, which lose TMEM119 expression^34,35^. As we could replicate both MS1 and MS2 microglia by experimentally inducing chronic meningeal inflammation *in vivo*, we propose that these MS-specific microglial phenotypes are the direct result of MS-related meningeal inflammation. Since CMI is induced by chronic overexpression of both TNFα and IFNγ in meninges of the sagittal sulcus, and TNFα and IFNγ are both extensively produced in inflamed MS meninges^38^, we assume that these cytokines are involved in driving the phenotypic changes in microglia. Whether the difference between MS1 and MS2 microglia is solely the result of a difference in duration of exposure to these cytokines or whether other (B cell derived) inflammatory factors play a role, needs to be further investigated.

Next to a direct effect of meningeal inflammation on microglia, our data also indicate that neurons act as intermediaries and actively attract microglia to their cell soma. Similar to findings under different neuroinflammatory conditions^23,30^, we show that the juxtaposition of microglia to neuronal somata leads to displacement of pre-synapses from the cell soma. At the same time, we observed increased synapse-phagocytosis by microglia. Interestingly, both displacement and phagocytosis of synapses was predominantly seen in MS1 patients. In contrast, significant neuronal loss was restricted to cortical layers 2 and 3 of MS2 cases, which is in line with a recent study showing a specific loss of CUX2^+^ neurons in the same layers, which spatially associated with inflamed meninges^29^. Given the differences in neuronal pathology between MS1 and MS2 cases, we speculate that pre-synaptic displacement and stripping by MS1 microglia could represent a protective response to neuroinflammation and prevent neuronal loss, as has previously been observed during LPS-induced neuroinflammation^23^. In contrast, MS2 microglia might have lost their ability to protect neurons from degenerating and perhaps even actively contribute to neuronal damage. If true, it would be important to investigate whether these dichotomous microglial phenotypes are mainly driven by differences in the neuroinflammatory environment, e.g. more meningeal B cells in MS2 cases, or the result of microglial exhaustion over time, as suggested by our *in vivo* data.

Surprisingly, we did not detect an overall decrease in pre-synaptic density in SPMS cortex, despite evidence for increased synapse phagocytosis by microglia. Although loss of synapses has been extensively reported in hippocampus^39,40^, thalamus^31^ and spinal cord of MS patients^41^, evidence for a loss of synapses in the cortex is more ambiguous. Reduced spine density on apical dendrites, indicating post-synaptic loss, has been shown in both myelinated and demyelinated cortex of MS patients^9^, whereas other studies, using similar methods to ours, only found a significant loss of synaptophysin in leukocortical demyelinated lesions but not in normal appearing MS cortex^42^ and no correlation with cortical atrophy^43^. Furthermore, the lack of decreased synaptic density in our cohort might be explained by the low rate of synapse removal (<0,5% of synapses within microglial lysosomes) in progressive MS cortex, which might allow for compensatory synaptogenesis, alternatively concomitant cortical thinning in these patients might obscure a reduction in synapse density despite an overall loss of synapses. Similarly, we also did not observe a significant difference in parameters of disease progression between the MS clusters, which one might expect given the differences in meningeal inflammation and neuronal loss and their strong correlation with disease severity^8,13^. However, this could be a limitation of our experimental setup, in which we only analyzed one cortical region in each patient. And given the strong spatial association between meningeal inflammation, microglial activation and neuronal damage^8,29^, pathology in this cortical region might not be predictive for the extent of tissue damage in the remaining cortex.

In this study, the results obtained from what we coined the CMI animal model, corroborates the initial experiments using this animal model^18^, including persistent meningeal inflammation, cortical demyelination and neuronal loss. Although our data indicate that MOG-immunization prior to lentiviral injection does not significantly alter the extent of meningeal inflammation, we did find that the difference between 1 month and 2 month microglia was exacerbated in MOG-immunized rats which is why we decided to focus on these animals. It would be interesting to explore if this difference is caused by increased cortical demyelination in MOG-immunized animals^18^, or by the presence of MOG-primed lymphocytes in the meninges. Taken together, we convincingly show that our experimental model of chronic meningeal inflammation closely mimics meningeal inflammation-induced cortical pathology in progressive MS patients. Although this model strongly emphasizes the role of TNFα and IFNγ in driving cortical pathology, and thereby likely over-simplifies the complex pathological processes in MS meninges and cortex, we are convinced that this model will be a valuable addition to our toolbox for studying progressive MS.

In conclusion, we have uncovered two distinct MS-specific microglial phenotypes in the cortex of progressive MS patients that are driven by local meningeal inflammation and differentially associate with neuronal damage. Results obtained in a novel experimental model for chronic MS-like meningeal inflammation suggest that these phenotypes occur sequentially and that microglia lose their protective properties over time, leading to neuronal loss. Hence, timely targeting of the processes contributing to microglial activation in the progressive MS cortex provides an interesting therapeutic strategy to combat progressive MS.

## Methods

### Human tissue samples

Post-mortem fixed-frozen blocks containing cingulate or insular cortex gyrus from 20 confirmed secondary progressive MS (SPMS) cases (mean age of death = 48.9, range 35-65) and 6 non-neuroinflammatory controls (mean age of death = 62, range 35-77) were provided by the UK MS Society Tissue Bank at Imperial College London. All donors or their next of kin provided fully informed consent for autopsy and use of material for research under ethical approval by the National Research Ethics Committee (08/MRE09/31). Relevant clinical and demographic information of individual control and SPMS cases are listed in Table 1. Tissue blocks were cut into 10 and 20 µm sections, and stored at -80 ºC until further use. In some analyzes, we had to omit 1 or 2 cases due to absence of meningeal tissue and upper cortical layers or loss of antigenicity in the corresponding tissue sections (listed in Table 1).

**Table 1:** Clinical and demographic data.

| Case ID | Gender (m/f) | Age at death (yr) | Post-mortem delay (hrs) | Cause of death | Disease duration (yr) | Time progressive (yr) | Time to wheel-chair (yr) |
| --- | --- | --- | --- | --- | --- | --- | --- |
| C14 | m | 64 | 18 | Cardiac failure | NA | NA | NA |
| C25 | m | 35 | 22 | Carcinoma of the tongue | NA | NA | NA |
| C28 | f | 60 | 13 | Ovarian cancer | NA | NA | NA |
| C36 | m | 68 | 30 | Heart failure, fibrosing alveolitis, arterome | NA | NA | NA |
| C45 | m | 77 | 22 |  | NA | NA | NA |
| C48<sup>1</sup> | m | 68 | 10 | Metastatic colon cancer | NA | NA | NA |
| MS330 | f | 59 | 21 |  | 39 | 24 | 21 |
| MS342 | f | 35 | 9 |  | 5 | 1 | 1 |
| MS352 | m | 43 | 26 |  | 18 | 14 | 10 |
| MS371 | m | 40 | 27 | Pneumonia | 16 | 7 | 4 |
| MS377 | f | 50 | 22 | Pneumonia, MS | 23 | 3 | NA |
| MS402 | m | 46 | 12 | Pneumonia, MS | 26 | 9 | 7 |
| MS403 | f | 54 | 11 | MS | 26 | 15 | 6 |
| MS404 | f | 55 | 17 | Septicaemia, shock, pneumonia | 34 | 20 | 2 |
| MS405 | m | 62 | 12 | Septicaemia, MS, metastatic colon cancer | 25 | 20 | 18 |
| MS408<sup>2</sup> | m | 39 | 21 | Pneumonia, sepsis | 11 | 5 | 2 |
| MS426<sup>3</sup> | f | 48 | 21 | MS | 19 | 19 | 13 |
| MS438 | f | 53 | 17 | MS | 12 | 4 | 4 |
| MS445<sup>4</sup> | f | 62 | 13 | Lower respiratory tract infection, MS | 38 | 13 | 15 |
| MS461 | m | 43 | 13 | Pneumonia, MS | 21 | 12 | 12 |
| MS473<sup>5</sup> | f | 39 | 9 | Pneumonia, MS | 13 | 10 | 6 |
| MS513 | m | 51 | 17 | MS, respiratory failure | 18 | 15 | 14 |
| MS527<sup>6</sup> | m | 47 | 10 | Pneumonia, MS | 25 | 15 | 13 |
| MS528 | f | 45 | 17 | MS | 25 | 10 | 3 |
| MS530<sup>7</sup> | m | 42 | 15 | MS | 21 | 15 | 14 |
| MS535<sup>8</sup> | f | 65 | 12 | MS | 40 | 25 | 20 |

<sup>1</sup> Excluded from Fig. 2f; Fig. 3b,f; Fig. 5e.
<sup>2</sup> Excluded from Fig. 3e.
<sup>3</sup> Excluded from Fig. 5e (all layers).
<sup>4</sup> Excluded from Fig. 2g.
<sup>5</sup> Excluded from Fig. 2f.
<sup>6</sup> Excluded from Fig. 2h.
<sup>7</sup> Excluded from Fig. 1c-f (layer 1).
<sup>8</sup> Excluded from Fig. 5e (all layers).

### Lentiviral vector production

Lentiviral vectors carrying human TNFα and IFNγ were produced as described previously^18^. In summary, HEK-293 cells were transfected with HIV-1 transfer plasmid (pRRL-sincppt-CMV-TNF-WPRE or pRRL-sincppt-CMV-IFN-WPRE genome plasmid), the packaging vector plasmids expressing HIV-1 gag/pol gene (pMD2-LgRRE), VSV-G envelope plasmids (pMD2-VSV-G) and HIV-1 Rev (pRSV-Rev) using CaCl_2_. Fresh medium supplemented with 10 mM sodium butyrate was added after 16h. The supernatant was harvested and filtered through a 0.45 μm filter after 36h, and centrifuged overnight. Lentiviral vectors were concentrated using ultracentrifugation and resuspended with TSSM (10 mM Tromethamine, 100 mM NaCl, 10 mg/mL sucrose and 10 mg/mL mannitol). The genome copy number was calculated using the Clontech Lenti-X qRT-PCR Titration kit (Takara Bio).

### Chronic meningeal inflammation animal model

8-10 week-old female Dark Agouti rats (140-160 g) were obtained from Janvier Labs (France). Rats were housed in groups of four in a 12h light/dark cycle and had ad libitum access to food and water. The UK Home Office approved all procedures. The induction of chronic meningeal inflammation (CMI) in Dark Agouti rats was performed as previously published^18^. Briefly, rats were anaesthetized with isofluorane and a sub-clinical autoimmune response to myelin was induced by injecting intra-dermally 5 µg recombinant mouse myelin oligodendrocyte glycoprotein (MOG; amino acids 1-119) diluted in phosphate buffered saline (PBS) and emulsified in incomplete Freund’s adjuvant (IFA, Sigma) or by injecting PBS alone emulsified in IFA as control (IFA ctrl). 20-23 days post-MOG or -IFA injection, rats underwent stereotactic surgery under isofluorane anaesthesia. A 5 mm hole was drilled in the skull in the midline 0.9 mm caudal to bregma. A calibrated glass capillary needle attached to a fixed-needle 10 μL Hamilton syringe was inserted to a depth of 2.3 mm down the sagittal sulcus, below the meningeal dura mater layer. The lentiviral vector mixture containing of 5 × 10^8^ genomic copies of TNFα and 5 × 10^7^ genomic copies of IFNγ was injected with a rate of 0.20 µL/mL. To allow diffusion of the mixture from the area of injection, the needle was left for 10 min in place and then slowly withdrawn. Under sodium pentobarbitone anaesthesia, rats were perfused 28 and 56 days after lentiviral injection with PBS followed by 4% paraformaldehyde (PFA). Brains were subsequently removed, post-fixed in 4% PFA overnight, cryoprotected in 30% sucrose and cut into 10 µm sections. Sections were stored at -80 ºC until further use. Naïve animals that did not receive any injection and MOG-immunized rats without stereotactic viral injection were used as controls.

### Immunolabeling

Sections were defrosted at room temperature and subjected to epitope retrieval at 95 ºC for 30 min followed by a blocking step at room temperature for 30 min with a solution containing 0.01% Triton X-100 (Sigma) and the appropriate 10% normal serum in PBS. Sections were incubated overnight at 4 ºC with primary antibodies (for details, see table 2) followed by a 2h incubation in Alexa fluorophore-labeled secondary antibodies (Thermo Fisher Scientific) for immunofluorescence. All immunofluorescent-stained sections were counterstained for DNA using DAPI (1:10000, Molecular Probes) followed by quenching of auto-fluorescence with 0,03% Sudan Black (Sigma) in 70% ethanol for 5 min. After washing, sections were embedded in Mowiol mounting medium and stored in the dark at 4 ºC until image acquisition.

**Table 2:** Primary antibody details.

| Antigen | Species | Used for | Dilution | Antigen Retrieval | Manufacturer | Cat. number |
| --- | --- | --- | --- | --- | --- | --- |
| CD19 | Rabbit | Human | 1:100 | EDTA-Tris pH 8 | Abcam | ab134114 |
| CD3 | Mouse | Human | 1:20 | EDTA-Tris pH 8 | DAKO | M7254 |
| CD3 | Mouse | Rat | 1:500 | without | BD Pharmigen | 550295 |
| CD4 | Rabbit | Human | 1:150 | Citrate pH 6 | Abcam | ab133616 |
| CD68 | Mouse | Human | 1:100 | Citrate pH 6 | Abcam | Ab995 |
| CD79a | Mouse | Rat | 1:200 | without | ThermoFisher | MA5-13212 |
| CD8 | Mouse | Human | 1:1000 | Citrate pH 6 | DAKO | M7103 |
| HLA class II (CR3/43) | Mouse | Human | 1:500 | Citrate pH 6 | DAKO | M077501-2 |
| HLA class II (OX6) | Mouse | Rat | 1:50 | without | In house |  |
| HuC/D | Mouse | Human/Rat | 1:1000 | Citrate pH 6 / EDTA-Tris pH 8 | ThermoFisher | A21271 |
| IBA1 | Goat | Human/Rat | 1:500 | Citrate pH 6 / EDTA-Tris pH 8 / without | Abcam | ab5076 |
| IBA1 | Rabbit | Human/Rat | 1:1000 | Citrate pH 6 / EDTA-Tris pH 8 / without | WAKO | 019-19741 |
| LAMP1 | Rabbit | Human | 1:500 | Citrate pH 6 | Cell Signaling | 9091P |
| LAMP1 | Rabbit | Rat | 1:1000 | EDTA-Tris pH 8 | Abcam | ab24710 |
| MOG | Mouse | Human/Rat | 1:50 | Citrate pH 6 | Reynolds Lab | N.A. |
| NeuN | Mouse | Rat | 1:1000 | Citrate pH 6 | Millipore | MAB377 |
| P2RY12 | Rabbit | Human | 1:100 | Citrate pH 6 | AnaSpec | 55042A |
| P2RY12 | Rabbit | Rat | 1:100 | without | AnaSpec | 55043A |
| Synaptophysin | Mouse | Human/Rat | 1:500 | Citrate pH 6 / EDTA-Tris pH 8 | DAKO | M7315 |
| TMEM119 | Rabbit | Human | 1:500 | Citrate pH 6 | Atlas abcam | HPA051870 |
| vGAT | Mouse | Rat | 1:500 | EDTA-Tris pH 8 | Synaptic Systems | 131 011 |

For immunohistochemistry, primary antibodies were visualized using the EnVision^+^ visualization system with 3,3’-diaminobenzidine (DAB) as the chromogen (DAKO). Sections were subsequently counterstained with hematoxylin, dehydrated, and embedded in Entellan medium (Merck). Sections were stored at room temperature until image acquisition.

### Image acquisition and analysis

All immunofluorescent stained sections were imaged on a Nikon A1R laser-scanning confocal microscope equipped with a resonant scanner, except sections stained for synaptic markers which were imaged on a Leica TCS SP8 confocal microscope equipped with a white light laser and HyD hybrid detectors. In human tissue, all confocal images were taken in the cortex surrounding the deeper part of a sulcus, whereas in rats all images were taken in the sagittal sulcus and the cortex directly adjacent. Acquisition details for individual datasets are described in the corresponding sections below. DAB stained sections were scanned on a Vectra Polaris whole-slide scanner (Akoya Biosciences) using either a 20x (MOG and NeuN) or 10x (HuC/D) objective.

#### Microglial morphology

Confocal images were acquired from cortical layers 1, 3 and 5/6 in 20 µm (human tissue) or 10 µm (rat tissue) IBA1-stained sections using a 40x objective and a z-stepsize of 1 µm (human) or 63x objective and a z-stepsize of 0.1 µm (rat). IBA1^+^ cells were manually traced from 2D maximum intensity projections of the aforementioned confocal images using FIJI (NIH). Per image, around 10-20 cells were selected for tracing. Cells were randomly selected, but had to meet several inclusion criteria: 1) cells should be completely included within the z-stack borders of the image; 2) cells should not overlap with one another; 3) cells should not be associated to a vessel. Next, traced microglia were analyzed using the Sholl Analysis Plugin^44^ with a 0.3 µm step size from the cell soma. Number of branches and junctions, and branch lengths were quantified by the AnalyzeSkeleton Plugin^45^ using the same microglia cell tracings. Soma surface area of the traced microglia was measured using the freehand selection tool in FIJI.

#### Microglial protein expression

Confocal images were acquired in cortical layer 3 of sections double-labeled for IBA1 together with P2Y12, TMEM119, HLA class II or CD68 using a 40x objective and a z-stepsize of 0.5 µm. Per section, two images were taken. Imaris software (Version 9.5.2, Bitplane AG) was used to quantify mean fluorescence intensity in microglia, by first segmenting all IBA1^+^ microglia using the surfaces function followed by measuring the total fluorescence intensity sum of the aforementioned markers in microglia divided by total microglial volume. Final mean fluorescence intensity was determined as the average of both images.

#### Meningeal immune-cells

3-6 confocal images were acquired in the meninges in and around sulci of sections stained for CD3/CD19, CD4/CD8 and IBA1 with a 20x objective and a z-step size of 1.4 µm. In rat sections stained for CD3/CD79a and IBA1, an image of the whole sagittal sulcus and its respective meninges was taken with a 20x objective and z-stepsize of 1 µm. The absolute number of positive cells and DAPI^+^ nuclei within the meninges were counted manually using FIJI. Cells and nuclei located in meningeal vessels were excluded. To determine meningeal immune cell ratios, absolute CD3^+^, CD19^+^/CD79a^+^ and IBA1^+^ cell numbers were normalized to the number of DAPI^+^ nuclei in the same sections. Human sections containing 10 or fewer positive immune cells in meningeal tissue were excluded from the immune cell ratio quantifications.

#### Demyelination and neuronal loss

In whole slide scans of MOG, NeuN and HuC/D stained sections, cortical layers were annotated using QuPath version v0.2.0-m7^46^. The annotated areas in MOG-stained sections were exported to FIJI, a manual threshold was set and the positive fraction per area was measured to determine MOG^+^ area in individual cortical layers. Secondly, in human MOG-stained sections, all completely demyelinated cortical lesions were outlined in FIJI and used to calculate the percentage of demyelinated cortex. Neuronal density was measured using the positive cell detection function in QuPath. DAB threshold was manually adjusted for each image to account for differences in staining intensity and all other parameters were kept the same for each subject.

#### Microglia-neuron association

Confocal images were acquired from cortical layers 3 and 5/6 in two different areas in sections co-stained for IBA1 and HuC/D with a 20x objective and z-stepsize of 1 µm. First, all HuC/D^+^ neuronal somata were marked in single-channel displays of each image using the Multi-Point tool in FIJI. The same tool was then used to mark all IBA1^+^ microglia cell bodies in the same images. The overlay image was used to count the amount of neuronal somata directly contacted by one or more microglial cell bodies, and vice versa. Only cell bodies containing a clearly visible DAPI^+^ nucleus were counted.

#### Peri-somatic synapse displacement

Confocal images were taken from cortical layers 3 (human and rat) and 5/6 (human) Synaptophysin, HuC/D and IBA1 labeled sections with a 100x objective (human) or 63x objective and 2.0 digital zoom (rat) and a z-stepsize of 0.1 µm. In single-plane confocal images of 6 MS patients and 3 CMI 1 and 2 month rats, HuC/D^+^ neuronal somata were outlined and their perimeter measured using FIJI. Next, neurons were classified as associated with IBA1^+^ microglia or not, and the length of the soma in contact with microglia was measured in FIJI. Lastly, all Synaptophysin^+^ pre-synaptic contacts on the outlined neuronal soma were measured and used to calculate the percentage of the neuronal soma that was covered by pre-synapses.

#### Synapse phagocytosis and synaptic density

15-30 confocal images were acquired from cortical layer 3 of stained sections for DAPI/IBA1/LAMP1/Synaptophysin (human) and DAPI/IBA1/LAMP1/vGAT (rat) with a 100x (human) or 63x and 2.0 digital zoom (rat) and a z-stepsize of 0.1 µm. For synaptic density and synaptic engulfment by IBA1^+^ cells, maximum intensity projections of 2 z-planes (human) or full z-stack images (rat) were used. Using Imaris software, surfaces marking nuclei (DAPI^+^), microglia (IBA1^+^) and lysosomes (LAMP1^+^) were created with individual settings per donor to account for differences in staining intensity. Next, the spots function was used to detect pre-synapses (Synaptophysin^+^ in human / vGAT^+^ in rat) using a spot diameter of 0.5 µm and including a quality filter which was individually set for each donor/animal. Pre-synaptic density was quantified by dividing the total number of detected spots by the total volume of the images. Microglial phagocytosis of pre-synapses was calculated by identifying spots that had a maximum distance of 0 µm to IBA1^+^ LAMP1^+^ surfaces and dividing their number to the total volume of the images or the total number of spots, whereas phagocytosis of pre-synapse by non-microglial cells was quantified by counting spots with a maximum distance of 0 to IBA1^-^ LAMP1^+^ surfaces.The same images were used to quantify microglial LAMP1 expression, which was done as described in the Microglial Protein Expression section above, although now the average of 15-30 images was used to calculate mean fluorescence intensity.

### Principal Component Analysis

To explore if combining parameters of microglia morphology, phenotype and density could reveal different MS subgroups, we performed a principle component analysis (PCA) with scaling of the parameters using the web-based MetaboAnalyst (http://www.metaboanalyst.ca)^47^. We included measures for microglia morphology and shape (AUC derived from the sholl analysis and soma size), microglia protein expression (P2Y12, TMEM119, CD68, HLA class II) and microglial density. For this analysis, data from all analyzed layers was combined if applicable. The few missing values were estimated using the median value of the group. To detect and characterize sub-populations of MS patients based on their microglia phenotype, K-means clustering (k = 3) was applied and differences in individual microglial parameters between the subgroups were corroborated with appropriate statistical testing.

### Statistics

All analyzes were done blinded to disease or experimental condition. Graphpad Prism 8.2.1 was used for all statistical tests. First, we used Shapiro-Wilk and F-test to test for normality and equality of variances, respectively, and appropriate tests were selected accordingly. For comparing two experimental groups, unpaired two-tailed Student’s t-test with or without Welch’s correction for unequal variances, or Mann-Whitney test was used. For comparing more than two groups, we used two-tailed one-way analysis of variance (ANOVA) with Tukey test for multiple comparisons, Welch ANOVA followed by Dunnett’s T3 multiple comparisons test, or Kruskal-Wallis test followed by Dunn’s multiple comparisons test. Data were judged to be statistically significant when p < 0.05 and, if significant, reported in the figures using the significance levels indicated in the figure legends.

## Acknowledgements

We thank the Microscopy and Cytometry Core Facility and Mike de Kok of the Amsterdam UMC for excellent technical support. We thank the Mo2Ab Core Facility for providing the HLA class II (OX6) antibody. We thank the UK MS Society Tissue Bank at Imperial College and Dr. Djordje Gveric for the supply of human post-mortem tissue samples. This work was supported by the National MS Society (grant RG-1601-07456 to RR and JvH) and the Dutch MS Research Foundation MS18-358 to HEdV.

## Author contributions

LvO and CRM performed experiments, analyzed data, contributed to design of the study and wrote the manuscript. CPM performed experiments and revised the manuscript. SK, REJ, AK, SMAvdP, LK, ED and MF performed experiments and analyzed data. GS, JJG, SA and NDM provided valuable scientific input and revised the manuscript. JvH and HEdV contributed to design of the study, obtained research grants and revised the manuscript. RR conceived and supervised the study, obtained research grants and revised the manuscript. MEW conceived and supervised the study, performed experiments, analyzed data and wrote the manuscript.

## Notes

### Competing Interest Statement

The authors have declared no competing interest.

## References

1. Compston, A. & Coles, A. Multiple sclerosis. The Lancet 372, 1502–1517 (2008).

2. Miller, D. H., Hornabrook, R. W. & Purdie, G. The natural history of multiple sclerosis: A regional study with some longitudinal data. J. Neurol. Neurosurg. Psychiatry 55, 341–346 (1992).

3. Lublin, F. D. et al. Defining the clinical course of multiple sclerosis: The 2013 revisions. Neurology 83, 278–286 (2014).

4. Hauser, S. L. & Oksenberg, J. R. The Neurobiology of Multiple Sclerosis: Genes, Inflammation, and Neurodegeneration. Neuron 52, 61–76 (2006).

5. Faissner, S., Plemel, J. R., Gold, R. & Yong, V. W. Progressive multiple sclerosis: from pathophysiology to therapeutic strategies. Nat. Rev. Drug Discov. (2019). doi: 10.1038/s41573-019-0035-2

6. Luchetti, S. et al. Progressive multiple sclerosis patients show substantial lesion activity that correlates with clinical disease severity and sex: a retrospective autopsy cohort analysis. Acta Neuropathol. 135, 511–528 (2018).

7. Peterson, J. W., Bö, L., Mörk, S., Chang, A. & Trapp, B. D. Transected neurites, apoptotic neurons, and reduced inflammation in cortical multiple sclerosis lesions. Ann. Neurol. 50, 389–400 (2001).

8. Magliozzi, R. et al. A Gradient of neuronal loss and meningeal inflammation in multiple sclerosis. Ann. Neurol. 68, 477–493 (2010).

9. Jürgens, T. et al. Reconstruction of single cortical projection neurons reveals primary spine loss in multiple sclerosis. Brain 139, 39–46 (2016).

10. Eijlers, A. J. C. et al. Predicting cognitive decline in multiple sclerosis: a 5-year follow-up study. doi: 10.1093/brain/awy202

11. Calabrese, M. et al. Cortical lesion load associates with progression of disability in multiple sclerosis. Brain 135, 2952–2961 (2012).

12. Calabrese, M. et al. Exploring the origins of grey matter damage in multiple sclerosis. Nat. Rev. Neurosci. 16, 147–158 (2015).

13. Howell, O. W. et al. Meningeal inflammation is widespread and linked to cortical pathology in multiple sclerosis. Brain 134, 2755–2771 (2011).

14. Serafini, B., Rosicarelli, B., Magliozzi, R., Stigliano, E. & Aloisi, F. Detection of Ectopic B-cell Follicles with Germinal Centers in the Meninges of Patients with Secondary Progressive Multiple Sclerosis. Brain Pathol. 14, 164–174 (2004).

15. Bell, L., Lenhart, A., Rosenwald, A., Monoranu, C. M. & Berberich-Siebelt, F. Lymphoid Aggregates in the CNS of Progressive Multiple Sclerosis Patients Lack Regulatory T Cells. Front. Immunol. 10, 1–18 (2020).

16. Gardner, C. et al. Cortical grey matter demyelination can be induced by elevated pro-inflammatory cytokines in the subarachnoid space of MOG-immunized rats. Brain 136, 3596–3608 (2013).

17. Magliozzi, R. et al. The CSF Profile Linked to Cortical Damage Predicts Multiple Sclerosis Activity. Ann. Neurol. (2020). doi: 10.1002/ana.25786

18. James, R. E. et al. Persistent elevation of intrathecal pro-inflammatory cytokines leads to multiple sclerosis-like cortical demyelination and neurodegeneration. Acta Neuropathol. Commun. 8, 66 (2020).

19. Nimmerjahn, A., Kirchhoff, F. & Helmchen, F. Resting microglial cells are highly dynamic surveillants of brain parenchyma *in vivo*. Science 308, 1314–8 (2005).

20. Sierra, A., Abiega, O., Shahraz, A. & Neumann, H. Janus-faced microglia: Beneficial and detrimental consequences of microglial phagocytosis. Frontiers in Cellular Neuroscience 7, 6 (2013).

21. Wu, Y., Dissing-Olesen, L., MacVicar, B. A. & Stevens, B. Microglia: Dynamic Mediators of Synapse Development and Plasticity. Trends in Immunology 36, 605–613 (2015).

22. Cserép, C. et al. Microglia monitor and protect neuronal function through specialized somatic purinergic junctions. Science (80-.). 367, 528–537 (2020).

23. Chen, Z. et al. Microglial displacement of inhibitory synapses provides neuroprotection in the adult brain. Nat. Commun. 5, 1–12 (2014).

24. Sharma, K., Wu, L. J. & Eyo, U. B. Calming Neurons with a Microglial Touch. Trends Neurosci. 43, 197–199 (2020).

25. Damisah, E. C. et al. Astrocytes and microglia play orchestrated roles and respect phagocytic territories during neuronal corpse removal *in vivo*. Sci. Adv. 6, 1–12 (2020).

26. Butovsky, O. & Weiner, H. L. Microglial signatures and their role in health and disease. Nat. Rev. Neurosci. 19, (2018).

27. Deczkowska, A. et al. Disease-Associated Microglia: A Universal Immune Sensor of Neurodegeneration. Cell 173, 1073–1081 (2018).

28. Magliozzi, R. et al. Meningeal B-cell follicles in secondary progressive multiple sclerosis associate with early onset of disease and severe cortical pathology. Brain 130, 1089–1104 (2007).

29. Schirmer, L. et al. Neuronal vulnerability and multilineage diversity in multiple sclerosis. Nature 1–8 doi: 10.1038/s41586-019-1404-z

30. Di Liberto, G. et al. Neurons under T Cell Attack Coordinate Phagocyte-Mediated Synaptic Stripping. Cell 1–14 (2018). doi: 10.1016/j.cell.2018.07.049

31. Werneburg, S. et al. Targeted Complement Inhibition at Synapses Prevents Microglial Synaptic Engulfment and Synapse Loss in Demyelinating Disease. Immunity 1–16 (2019). doi: 10.1016/j.immuni.2019.12.004

32. Mahad, D. H., Trapp, B. D. & Lassmann, H. Pathological mechanisms in progressive multiple sclerosis. Lancet Neurol. 14, 183–193 (2015).

33. Reali, C. et al. B cell rich meningeal inflammation associates with increased spinal cord pathology in multiple sclerosis. Brain Pathol. (2020). doi: 10.1111/bpa.12841

34. Krasemann, S. et al. The TREM2-APOE Pathway Drives the Transcriptional Phenotype of Dysfunctional Microglia in Neurodegenerative Diseases Article The TREM2-APOE Pathway Drives the Transcriptional Phenotype of Dysfunctional Microglia in Neurodegenerative Diseases. Immunity 47, 566-581.e9 (2017).

35. Keren-Shaul, H. et al. A Unique Microglia Type Associated with Restricting Development of Alzheimer’s Disease. Cell 169, 1276-1290. e17 (2017).

36. McGeer, P. L., Itagaki, S., Tago, H. & McGeer, E. G. Reactive microglia in patients with senile dementia of the Alzheimer type are positive for the histocompatibility glycoprotein HLA-DR. Neurosci. Lett. 79, 195–200 (1987).

37. Vogel, D. Y. S. et al. Macrophages in inflammatory multiple sclerosis lesions have an intermediate activation status. J. Neuroinflammation 10, (2013).

38. Gardner, C. et al. Cortical grey matter demyelination can be induced by elevated pro-inflammatory cytokines in the subarachnoid space of MOG-immunized rats. Brain 136, 3596–3608 (2013).

39. Dutta, R. et al. Demyelination causes synaptic alterations in hippocampi from multiple sclerosis patients. Ann. Neurol. 69, 445–454 (2011).

40. Michailidou, I. et al. Complement C1q-C3-associated synaptic changes in multiple sclerosis hippocampus. Ann. Neurol. 77, 1007–1026 (2015).

41. Petrova, N. et al. Synaptic Loss in Multiple Sclerosis Spinal Cord. Ann. Neurol. 1–7 (2020). doi: 10.1002/ana.25835

42. Wegner, C., Esiri, M. M., Chance, S. A., Palace, J. & Matthews, P. M. Neocortical neuronal, synaptic, and glial loss in multiple sclerosis. Neurology 67, 960–967 (2006).

43. Popescu, V. et al. What drives MRI-measured cortical atrophy in multiple sclerosis? Mult. Scler. 1–11 (2015). doi: 10.1177/1352458514562440

44. Ferreira, T., Ou, Y., Li, S., Giniger, E. & van Meyel, D. J. Dendrite architecture organized by transcriptional control of the F-actin nucleator Spire. Dev. 141, 650–660 (2014).

45. Arganda-Carreras, I., Fernández-González, R., Muñoz-Barrutia, A. & Ortiz-De-Solorzano, C. 3D reconstruction of histological sections: Application to mammary gland tissue. Microsc. Res. Tech. 73, 1019–1029 (2010).

46. Bankhead, P. et al. QuPath: Open source software for digital pathology image analysis. Sci. Rep. 7, (2017).

47. Pang, Z., Chong, J., Li, S. & Xia, J. Metaboanalystr 3.0: Toward an optimized workflow for global metabolomics. Metabolites 10, (2020).

