## Supplementary Data for "Meningeal inflammation in multiple sclerosis induces phenotypic changes in cortical microglia that differentially associate with neurodegeneration"

### Supplementary figures

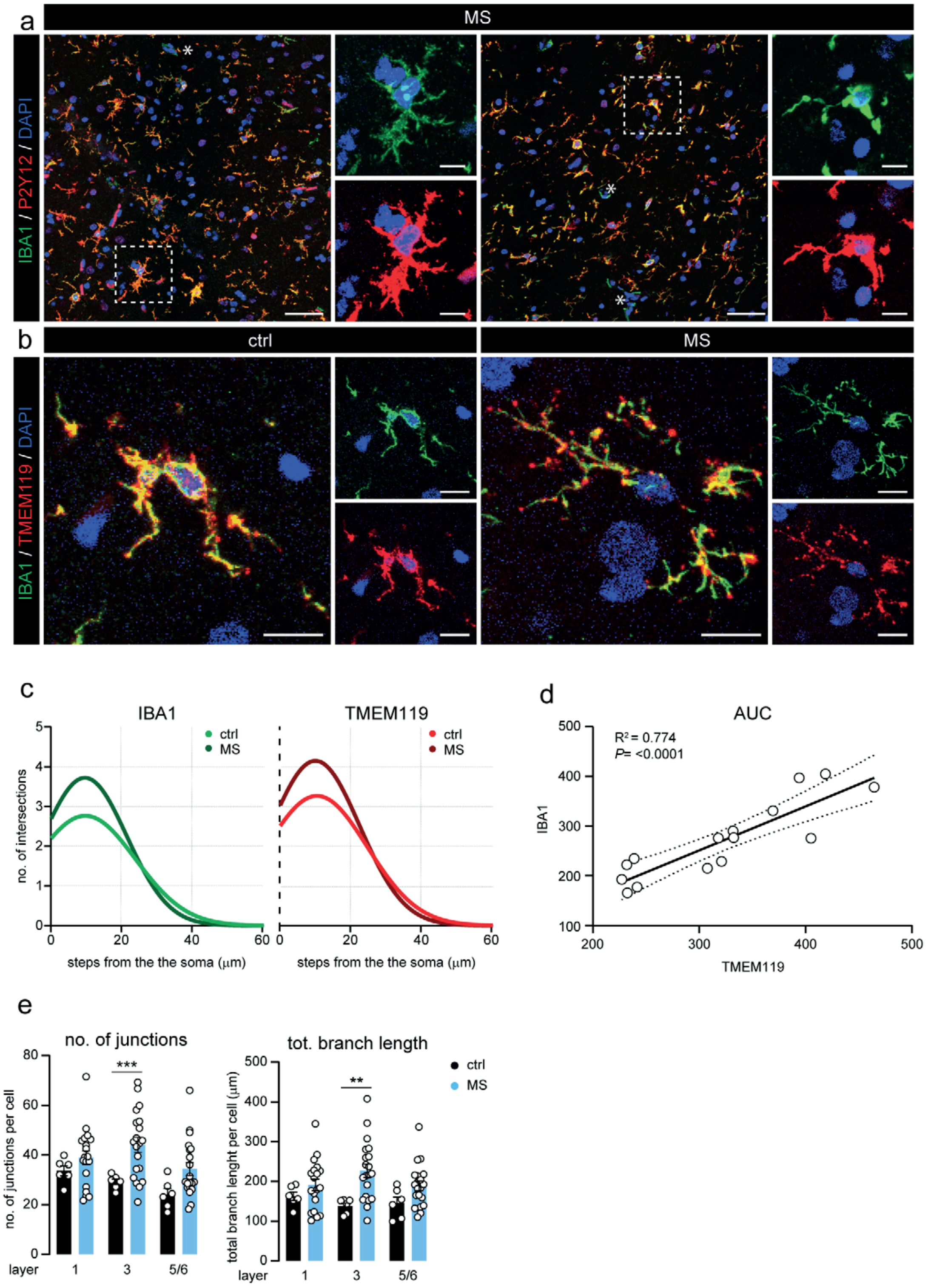


**Suppl. Fig. 1 | a.** Representative image of P2Y12 and IBA1 co-expression in cortical layer 3 of a ctrl and MS subject. Asterisks depict IBA1$^+$ vessel-associated cells that are negative for P2Y12 expression. **b.** Micrographs of single IBA1$^+$, TMEM119$^+$ cortical microglia in layer 3 from a ctrl and MS patient. **c**. Non-linear curve fit of the average number of microglial branch intersections per 0.3 µm step from the cell soma per cortical layer of IBA1$^+$ and TMEM119$^+$ cells as measured by Sholl analysis. **d**. Quantification of the correlation between the total Sholl-derived area-under-the-curve (AUC) of IBA1$^+$ and TMEM119$^+$ cells. **e.** Quantification of the number of junctions and total branch length of microglia in the different neuronal layers of microglial cell morphology. Individual datapoints indicate averaged data from an individual donor, columns and error bars show mean ± SEM; **p < 0.01, ***p < 0.001; *n* = 6 ctrl and *n* = 9 MS **(c, d)**, *n* = 6 ctrl and *n* = 20 MS **(e)**; Scale bars = 25 µm (**a**; overview), 10 µm (**a**: close-up), 20 µm **(b)**.

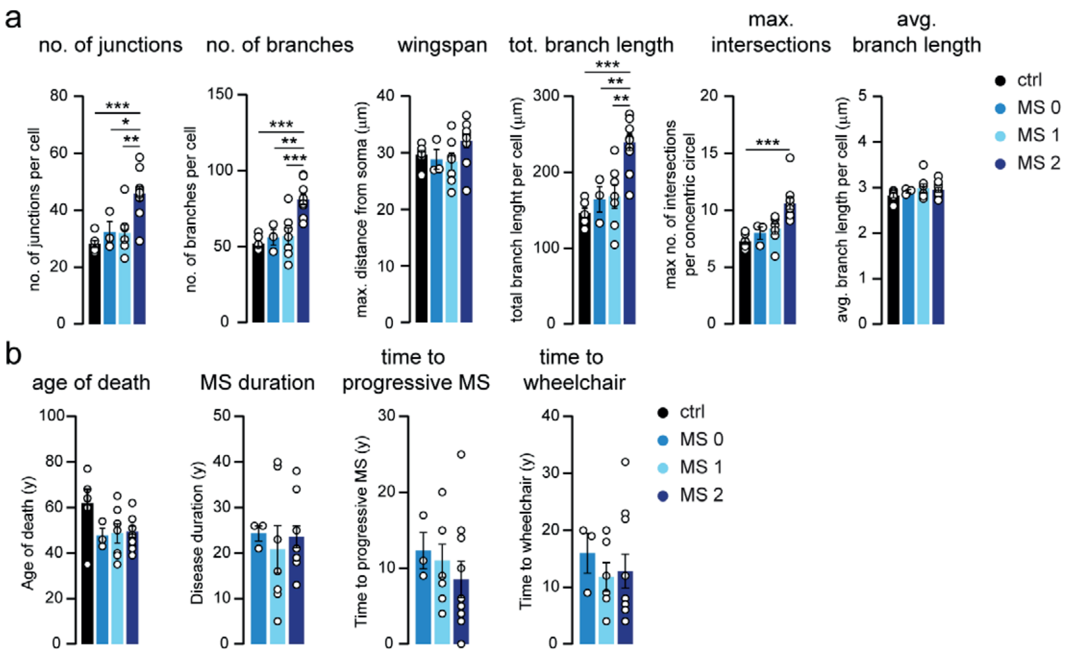


**Suppl. Fig. 2 | a.** Quantification of the average number of junctions, number of branches, wingspan, total branch length, maximum number of intersections and average branch length of microglial cell morphology in the different MS sub-clusters as detected by K-means clustering. **b.** Quantification of the age of death, MS disease duration, time from disease onset to progressive MS and time from disease onset to wheelchair bound in the different MS sub-clusters as detected by K-means clustering. Individual datapoints indicate averaged data from an individual donor, columns and error bars show mean ± SEM; *p < 0.05, **p < 0.01, ***p < 0.001; *n* = 6 ctrl, *n* = 3 MS0, *n* = 7 MS1, *n* = 10 MS2.

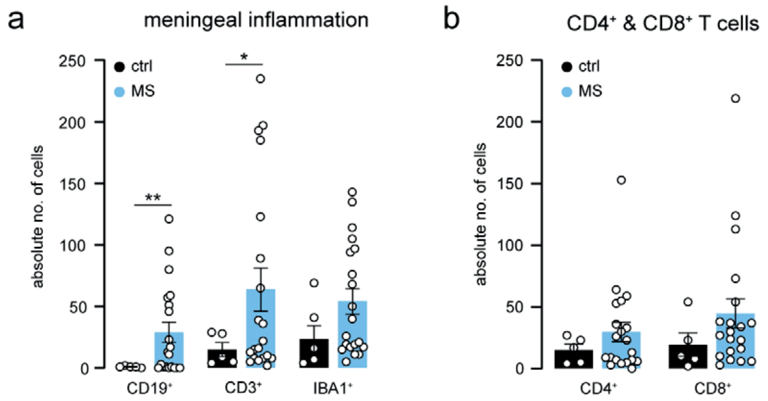


**Suppl. Fig. 3 | a.** Quantification of the absolute cell count of $CD19^+$ B cells, $CD3^+$ T cells and $IBA1^+$ myeloid cells in the meninges of ctrl and MS cases. **b**. Quantification of the absolute cell count of $CD4^+$ and $CD8^+$ T cells in the meninges of ctrl and MS subjects. Individual datapoints indicate averaged data from an individual donor, columns and error bars show mean ± SEM; *p < 0.05, **p < 0.01; *n* = 5 ctrl, *n* = 20 MS; Scale bars = 20 µm.

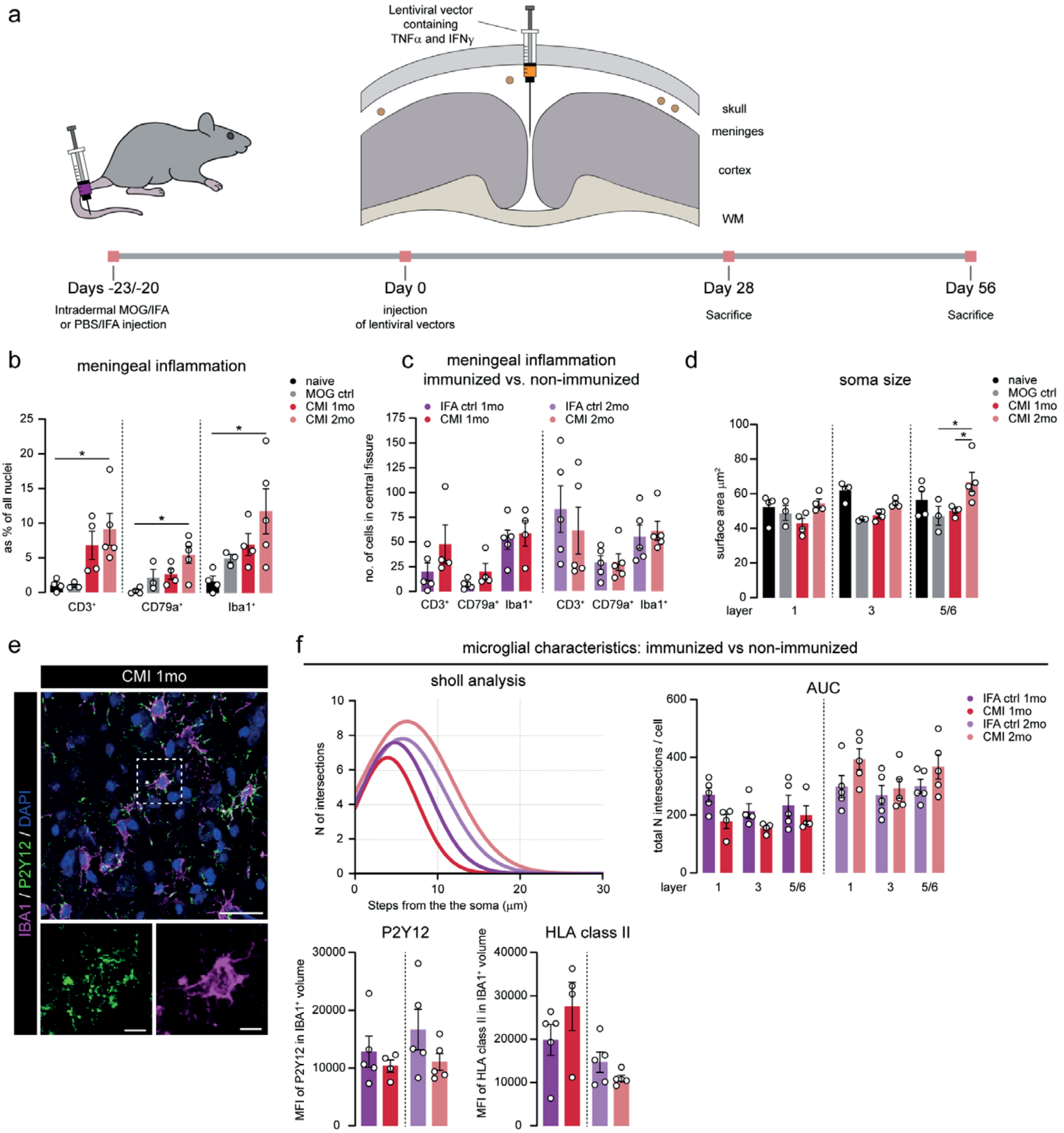


**Suppl. Fig. 4 | a.** Experimental setup of chronically induced meningeal inflammation in rats. Animals were either immunized with MOG/IFA (CMI) or injected with PBS/IFA (IFA ctrl) 23/20 days before injection of lentiviral constructs carrying the TNFα and IFNγ genes vectors in the sagittal sulcus. Animals were sacrificed at day 28 (1 month) or day 56 (2 months). **b.** Quantification of $CD3^+$ T cells, $CD79a^+$ B cells and $IBA1^+$ myeloid cells as percentage of all extravascular nuclei in the sagittal sulcus of naive, MOG ctrl, CMI 1 month and CMI 2 month animals. **c.** Absolute number of meningeal $CD3^+$, $CD79a^+$ and $IBA1^+$ cells in immunized (CMI) vs. non-immunized (IFA ctrl) rats. **d.** Quantification of the microglial soma size in layer 1, 3 and 5/6 of the cortex from naive, MOG ctrl, CMI 1 month and CMI 2 month animals. **e.** Representative image of cortical layer 3 of a CMI 1 month rat with the highest microglial density immunostained for IBA1 and P2Y12. **f.** Different measurements of $IBA1^+$ microglia: Non-linear curve fit of the average number of microglial branch intersections per 0.3 µm step from the cell soma as measured by Sholl analysis; total Sholl-derived area-under-the-curve (AUC); mean fluorescence intensity of P2Y12 and HLA class II in immunized (CMI) vs. non-immunized (IFA ctrl) rats. Individual datapoints indicate averaged data from an individual donor, columns and error bars show mean ± SEM; $*p < 0.05$, $**p < 0.01$, $***p < 0.001$; *n* = 4 naïve, *n* = 3 MOG ctrl, *n* = 5 IFA ctrl 1mo, *n* = 4 CMI 1mo, *n* = 5 IFA ctrl 2mo, *n* = 5 CMI 2mo **(b, c, e, f)**. Scale bars = 25 µm (**d**: overview); 5 µm (**d**: close-up).

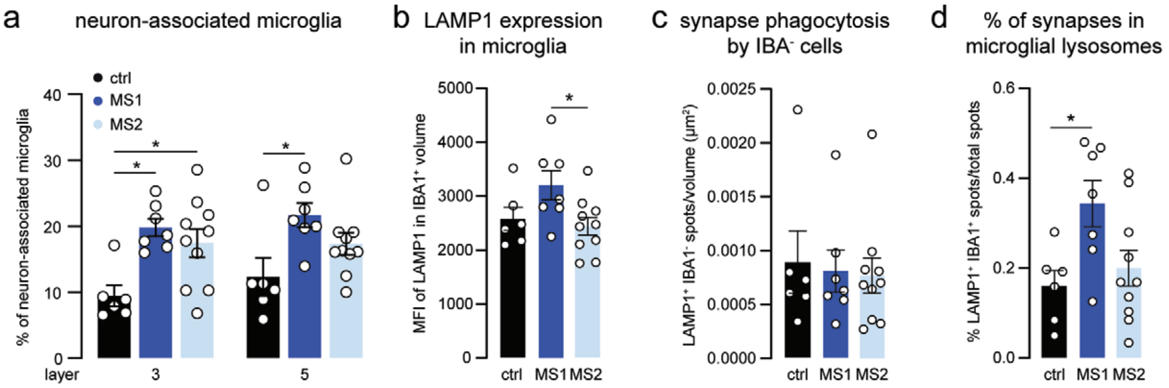


**Suppl. Fig. 5 | a.** Percentage of microglial cell bodies that are directly in contact with neuronal somata in cortical layers 3 and 5/6 of ctrl, MS1 and MS2. **b.** Quantification of the mean fluorescence intensity of LAMP1 in IBA1$^{+}$ volume in cortical layer 3 of ctrl, MS1 and MS2 patients. **c.** Density of synapses found within lysosomes of IBA1$^{-}$ cells in layer 3 of ctrl, MS1 and MS2 cases. **d.** Percentage of all Synaptophysin$^{+}$ synapses that are located in microglial lysosomes in cortical layer 3 of ctrl, MS1 and MS2. Individual datapoints indicate averaged data from an individual donor, columns and error bars show mean ± SEM; *$p < 0.05$; $n$ = 6 ctrl, $n$ = 7 MS1, $n$ = 10 MS2.

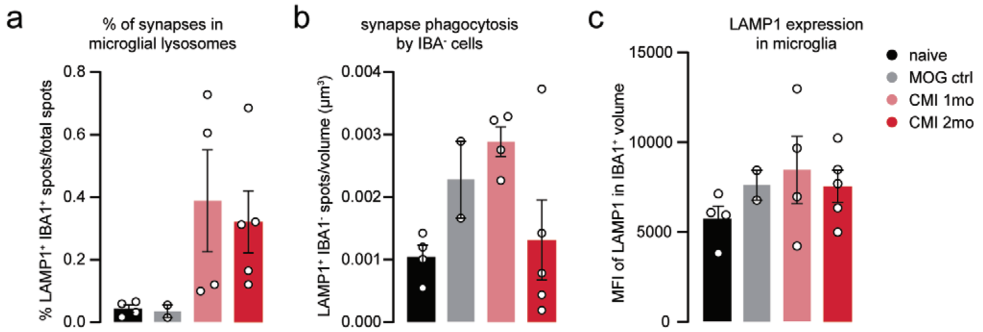


**Suppl. Fig. 6 | a.** Percentage of all vGAT⁺ synapses that are located in microglial lysosomes in cortical layer 3 of naive, MOG ctrl, CMI 1 month and CMI 2 month animals. **b.** Density of synapses found within lysosomes of IBA1⁻ cells in layer 3 of naive, MOG ctrl, CMI 1 month and CMI 2 month animals. **c.** Quantification of the mean fluorescence intensity of LAMP1 in cortical layer 3 microglia in naive, MOG ctrl, CMI 1 month and CMI 2 month animals. Individual datapoints indicate averaged data from an individual animal, columns and error bars show mean ± SEM; *n* = 4 naïve, *n* = 2 MOG ctrl, *n* = 4 CMI 1mo, *n* = 5 CMI 2mo.
